# Prefrontal PV interneurons facilitate attention and are linked to attentional dysfunction in a mouse model of absence epilepsy

**DOI:** 10.1101/2022.03.17.484733

**Authors:** Brielle R. Ferguson, John R. Huguenard

**Author notes:** ^*^Address for correspondence: John Huguenard, Ph.D., Department of Neurology, Stanford Neurosciences Building, 290 Jane Stanford Way, Stanford, Ca 94305, Correspondence: John Huguenard.

## Abstract

Absence seizures are characterized by brief periods of unconsciousness accompanied by a lapse in motor function that can occur hundreds of times throughout the day. Outside of these frequent moments of unconsciousness, approximately a third of patients experience treatment-resistant attention impairments. Convergent evidence suggests prefrontal cortex (PFC) dysfunction may underlie attention impairments in affected patients. To test this, we use a combination of slice physiology, fiber photometry, electrocorticography (ECoG), optogenetics, and behavior in the *Scn8a*^+/−^ mouse model of absence epilepsy. In these mice, we find decreased parvalbumin interneuron (PVIN) recruitment in the medial PFC (mPFC) *in vitro* and hypoactivity during cue presentation *in vivo* that is linked to attention dysfunction. Further, we find that low levels of mPFC PVIN activity are predictive of poorer performance in WT littermates. This highlights cue-evoked PVIN activity as an important mechanism for attention and suggests PVINs may represent a therapeutic target for cognitive comorbidities in absence epilepsy.

## Introduction

Attention is required for most aspects of moving through the world, from holding a conversation in a crowded room, to crossing a busy intersection. Disrupted attention presents as a persistent comorbidity across several disease states, even when certain hallmark symptoms respond to treatments or are in remission (Walter Heinrichs 2005; Rock et al. 2014). One such example is absence epilepsy, where attention deficits do not respond to current treatments and are a significant predictor of adverse long-term outcome across patients (Masur et al. 2013). Absence seizures are diagnosed by a distinct EEG pattern: bilateral synchronous 3-4 Hz spike-and-wave discharge (SWD), which manifests as brief periods of unconsciousness accompanied by a lapse in motor function (Gibbs et al. 1935, Panayiotopoulos 1999). Typical absence seizures are present in 10-15% of all epilepsy syndromes, and can occur hundreds of times throughout the day (Berg et al. 1999, Panayiotopoulos 2001, Posner 2008). While animal models reliably capture many aspects of the absence seizure phenotype (Vergnes et al. 1982, Coenen et al. 1987, Meeren et al. 2005, Papale et al. 2009), there has been little direct exploration of attention performance in rodent models of absence epilepsy or of the underlying mechanisms.

Components of attention include the initiation or engagement of attention, vigilance or sustained attention, and attentional shifting or flexibility (Mirsky 1987). Several studies point to the dorsolateral PFC supporting many of these components in humans and primates (Barcelo et al. 2000, Rossi et al. 2007, Zanto et al. 2011, Gregoriou et al. 2014). This is strengthened by lesion data from rodents of the closest functional homolog of human dorsolateral PFC, the prelimbic medial PFC (mPFC) (Muir et al. 1996, Chudasama et al. 2001, Kahn et al. 2012). Recent advances have allowed for more exploration into cell-type specific mechanisms of attention, revealing that fast-spiking GABAergic interneurons, or parvalbumin-expressing interneurons (PVINs), show attention-related activity. Medial PFC PVINs have been shown to represent rule information in a flexible attentional shifting task (Rikhye et al. 2018), while population increases in PVIN activity during the delay period prior to cue onset are associated with correct choices in a sustained attention task (Kim et al. 2016).

Here, we utilized *in vitro* local field potential recordings in mPFC slices, and *in vivo* fiber photometry, ECoG recordings, optogenetics, and behavior to explore circuit mechanisms of both normal attention and its disruption in the *Scn8a* mutant mouse model of absence epilepsy. In wild-type (WT) and *Scn8a* mutant mice, we observed that cue-evoked increases in mPFC PVIN activity were correlated with and predictive of correct choices during a novel attentional engagement task (AET). *Scn8a* mutant mice however, exhibited attenuated cue-evoked PVIN activity, reduced gamma band activity, and poorer performance on the AET. This was associated with reduced recruitment of mPFC feedforward inhibition in slice recordings. Finally, we found that gamma frequency optogenetic activation of PVINs during the cue period was sufficient to recover attention performance in *Scn8a* mutant mice. This suggests that cue-evoked increases in mPFC PVIN activity support basic attentional processes and may provide both an anatomical and cell-type specific therapeutic target for attention dysfunction in absence epilepsy.

## Results

### Scn8amutant mice exhibit attention deficits on the Attentional Engagement Task

To examine the mechanisms of attention impairments in absence epilepsy, we used mice with a heterozygous loss-of-function mutation in *Scn8a (Scn8a^med^)* (Kohrman et al. 1996) henceforward referred to as *Scn8a*^+/−^. Altered expression of *Scn8a* has been observed in humans with absence seizures and cognitive impairments (Trudeau et al. 2006, Berghuis et al. 2015). *Scn8a* encodes the voltage-gated sodium channel, NaV1.6, and mice with reduced NaV1.6 expression have frequent and well-characterized absence seizures, due at least in part to specific hypofunction of inhibitory neurons in the thalamus (Papale et al. 2009, Makinson et al. 2017). Still, the implications of partial *Scn8a* loss for cortical circuit function, including the mPFC and its role attentional processing, remain unexplored. To measure attention, we developed a novel assay, combining training and testing strategies from previous studies (Kahn et al. 2012, Turner et al. 2015, Wimmer et al. 2015). Mice (*Scn8a*^+/−^ and *Scn8a*^+/+^ littermates referred to as WT) were food-restricted to 85% of their free-feeding weight and habituated to the operant chamber. Using behavioral shaping, mice were trained to nose-poke in two distal ports to obtain a food reward (10 μL of condensed milk). Then, mice learned to initiate trials by nose-poking in a center port, and that a visual cue, a white LED present for 5 seconds (s) in one of the two reward ports, would indicate the location of a food reward (Fig. 1A). On average, mice learned this association in 6 training sessions, and there were no group differences in how long it took mice to reach stable performance or criterion (3 days at greater than 70 % correct or 2 days greater than 90% correct, Fig. 1A, bottom). Then, testing began, and while the cue still indicated the reward location, the cue length was varied pseudorandomly between 5s, 2s, 1s, 500ms, or 100ms. We chose this range of cue lengths, hypothesizing that the long 5s cue would require little attention due to the ease of detectability and prior exposure to this cue length during training. By contrast, the shorter cue lengths would be unexpected and significantly more difficult to report correctly, thus increasing cognitive and attentional load (Parasuraman et al. 1987). Additionally, by including the shortest cue lengths, 500ms and 100ms, we could approach a floor effect where most mice would have difficulty detecting the cue and we would expect them to perform poorly. With this design, we could identify specific intermediate cue lengths that would generate variable performance rates across mice and genotypes (Dinstein et al. 2015). We also hypothesized that performance at these intermediate cue lengths would be accompanied by related functional differences in brain activity (Faisal et al. 2008). Further, to isolate basic processes related to attentional recruitment, referred to here as engagement, rather than vigilance or selective attention such as is described elsewhere (Young et al. 2009, St Peters et al. 2011), we did by not incorporate any delays, distractors, or abstraction. In all mice, we observed a performance decrement with cue length, approaching chance levels at the 100ms cue. However, *Scn8a*^+/−^ mice performed significantly worse than WT mice at intermediate cue lengths (2s and 1s), suggesting an impairment in attentional engagement (Fig. 1B).

**Figure 1.**
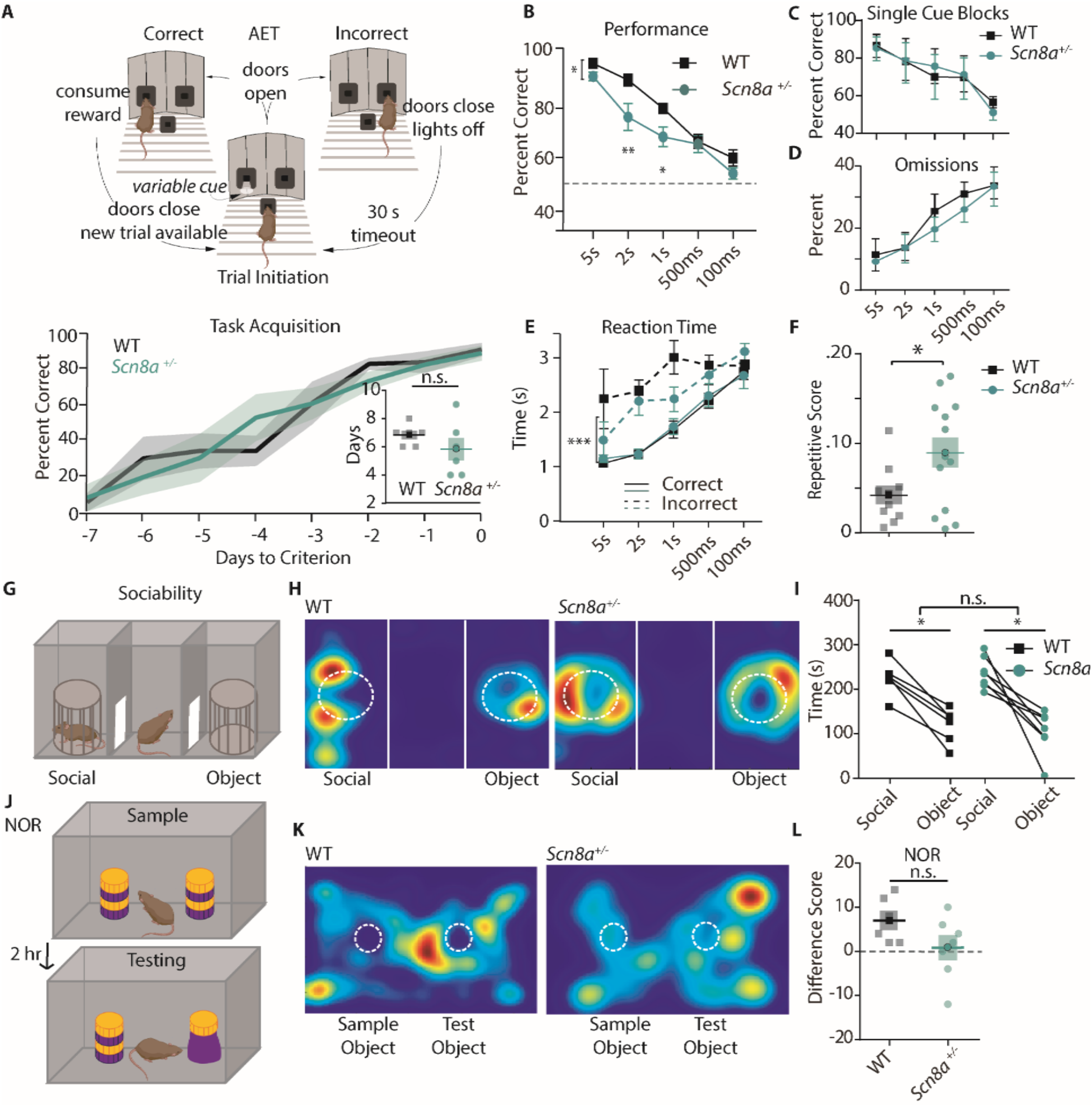
Scn8a^+/−^ mice exhibit attention deficits on the Attentional Engagement Task (AET). A, Top, AET task design. Bottom, Training accuracy for mice learning the AET. Lines represent performance displayed as average ± SE until mice reached criterion (WT are in black and *Scn8a*^+/−^ are in teal for this and all subsequent figures). Inset, comparison of total days to criterion, which was not different between groups (Student’s t test, p = 0.2666, n = 7 for both groups). B, *Scn8a*^+/−^ mice have reduced accuracy at 2s (p = 0.0049) and 1s (p = 0.0497) compared to WT littermates (two-way ANOVA, F_(1,70)_ = 16.17, p = 0.0001, n = 8 for both WT and *Scn8a*^+/−^). C, Animals were evaluated on their performance at each cue length in separate trial blocks, and there was no effect of genotype on performance in each session (F _(1, 40)_ = 0.0007687, p = 0.9780, n = 5 for both WT and *Scn8a*^+/−^). D, Omissions were not different between groups (two-way ANOVA, F_(1,70)_ = 0.8456, p = 0.3609, n = 8 for both WT and *Scn8a*^+/−^). E, There was no effect of group on reaction time, but there was an effect of accuracy (three-way ANOVA, group, F_(1,70)_ = 1.01, p = 0.3161; accuracy, F_(1,70)_ = 67.66, p = 0.0001;). F, The repetitive score was increased in *Scn8a*^+/−^ mice (Student’s t test, p = 0.0381, n = 10 and 13 for WT and *Scn8a*^+/−^ respectively). G, Top, design of the three-chamber sociability task. H and K, Representative heatmap images. I, Each group showed a significant preference for the novel mouse versus the object (p < 0.0001), but there were no differences between groups (F _(1, 11)_ = 0.5648, n = 6 and 7 for WT and *Scn8a*^+/−^ respectively). J, Novel object recognition paradigm. L, there were no significant differences in the difference score (Student’s t test, p = 0.111, n = 6 and 7 for WT and *Scn8a*^+/−^ respectively). Data are shown as averages from individual animals or groups, and errors bars or shading represent ± SE in this and all future figures unless stated otherwise.

Importantly, when we kept the cue length constant within trial blocks, presumably reducing attentional load, there were no differences in accuracy between groups (Fig. 1C). We also quantified omissions, trials in which the animal took longer than 5s following cue termination to make a choice (likely because the animals failed to perceive the presence of the cue), and found no differences between groups (Fig. 1D). This lack of difference in performance during single cue-length blocks or omissions strongly suggests that observed differences on the AET are specific to increases in attentional load rather than an effect of the loss of *Scn8a* on visual perception or discrimination. There was no effect of group on reaction time (seconds between cue termination and port selection), however all mice had increased reaction time for incorrect trials, which has also been correlated with impaired attention in other paradigms (Weissberg et al. 1990, Fitzpatrick et al. 2017) (Fig. 1E). To explore whether mice showed signs of behavioral inflexibility in their responses, we quantified repetitive responding, defined here as the number of blocks of 5 consecutive trials in which the animal selected the same port. We observed that *Scn8a*^+/−^ mice had a higher repetitive responding in comparison to WT littermates (Fig. 1F). Finally, to examine whether *Scn8a*^+/−^ mice exhibited broad behavioral impairments, we assayed sociability utilizing the three-chamber task and the hippocampal-dependent object recognition task, and observed no differences between groups (Fig. 1G-L). For subsequent AET experiments we utilized only three cue lengths in our analysis, 5s as the long cue, 2s as our intermediate (int) attention-related cue, and 500ms as the short cue to simplify the task interpretation, data presentation, and reduce the number of statistical comparisons.

### AET performance deficits are unrelated to seizure activity in Scn8a^+/−^ mice

Absence epilepsy patients experience attention impairments during interictal periods and while on successful treatment with anti-epileptic drugs (Barone et al. 2020), suggesting that attention dysfunction is not purely due to the periodic losses of consciousness associated with SWDs.

Additionally, absence seizure prevalence is highest in periods of quiet waking (Barone et al. 2020), thus we hypothesized that seizure burden would be low while animals were actively performing the task, and would not influence task performance. To examine this, we implanted *Scn8a*^+/−^ mice with screws for cortical surface (ECoG) recordings over the somatosensory cortex (S1) and recorded ECoG signals while animals were engaged in the task (Fig. 2A, B) or in their home cage. Engaging in the task reduced seizure frequency in *Scn8a*^+/−^ mice in comparison to that recorded in their home cage (Fig. 2C). To assess whether SWDs may contribute to reduced performance in *Scn8a*^+/−^ mice, we compared the ECoG spectral power using wavelet decomposition (Torrence et al. 1998), between correct trials and incorrect trials (Fig. 2D,E) in the Ø∗ (7-10 Hz) or β (10-20Hz) range (Fig. 2F,G) which encompass the fundamental and 2^nd^ harmonic frequency bands for absence seizures (Sorokin et al. 2017) (Fig. 2H). In *Scn8a*^+/−^ mice, there were no differences in power across cue lengths (Fig 2I-K). As we would expect a spectral power *increase* with any seizure activity (example SWD, Fig. 2H), the lack of difference in seizure band power during incorrect trials suggests that seizures do not contribute to AET errors in *Scn8a*^+/−^ mice. On average, in *Scn8a*^+/−^ mice, only ~1% of trials had seizures within 10 seconds of the trial, and in less than 1% of trials (0.24%) were we able to detect seizures during the cue (Fig 2-Supp. 1). Finally, power in seizure-related frequency bands was not different between *Scn8a*^+/−^ mice and WTs for the intermediate cue (Fig. 2L). This indicates that SWDs are largely absent while animals perform the AET, and differences in absence seizure activity do not explain reductions in performance in *Scn8a*^+/−^ mice.

**Figure 2.**
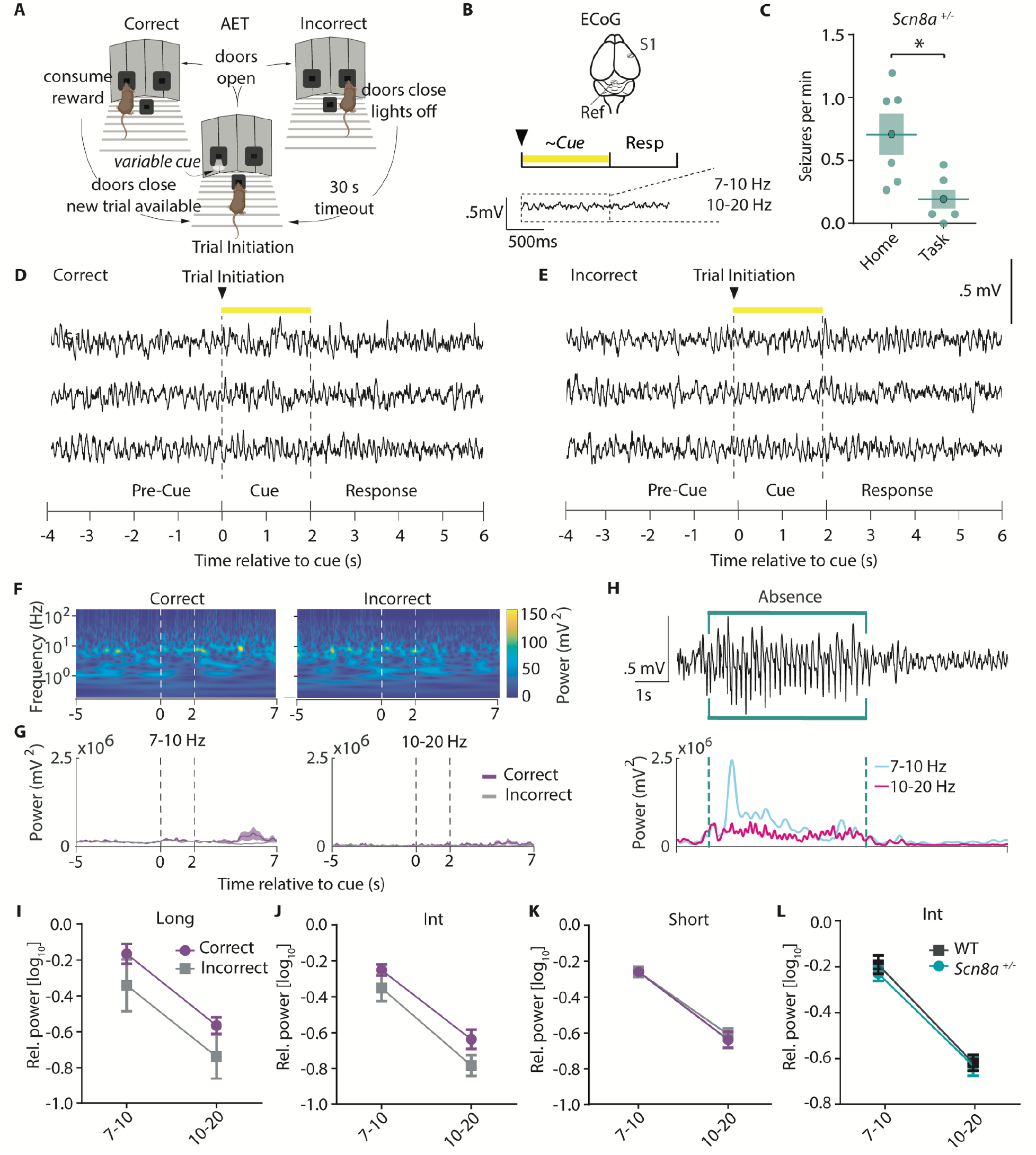
AET performance deficits are unrelated to seizure activity in Scn8a^+/−^ mice. A, AET design. B, Illustration of ECoG recording and metrics for power analysis. C, Seizures were reduced during the task (Student’s T test, p = 0.0155, n = 6 and 6 for home cage versus task animals respectively). D-E, Example ECoG activity from the highly seizure-active somatosensory cortex (S1) recorded during correct and incorrect trials with a 2s cue length. F, Representative time-frequency wavelet decomposition for correct trials (left) and incorrect trials (right). G, Trial-averaged power and variance over time in the Ø∗ (7-10 Hz) or β (10-20Hz) range with correct trials in purple and incorrect trials in grey. H, An example of a detected absence seizure (top) and the corresponding power in the Ø∗ (7-10 Hz) or β (10-20Hz) range. I-K, In *Scn8a*^+/−^ mice, there were no differences in power across cue lengths in the Ø∗ (7-10 Hz) or β (10-20Hz) range between correct and incorrect trials (I, Long cue, two-way ANOVA, accuracy effect, F _(1, 16)_ = 3.998, p = 0.0628; J, Int cue, two-way RM ANOVA, accuracy effect, F _(1, 10)_ = 3.367, p = 0.0964; K, short cue, two-way RM ANOVA, accuracy effect, F (1, 10) = 0.2082, p = 0.6580; n = 6 for all) L, There was no difference in average power between WT and *Scn8a*^+/−^ mice in the Ø∗ (7-10 Hz) or β (10-20Hz) range (two-way ANOVA, group effect, F _(1, 18)_ = 0.1185, p = 0.5101, n = 5 and 6 for WT and *Scn8a*^+/−^ mice respectively). See also Figure 2 Figure Supplement (Supp.) 1.

### Cue detection in the AET is mPFC-dependent and linked to attentional load

Given that our task is a novel adaptation of previous cue-based attention tasks, we wanted to explore whether the AET relied upon the mPFC, an area classically linked to attention across multiple species

(Ungerleider 2000, Chudasama et al. 2001, Everling et al. 2002). To do so, we implanted optical fibers targeting the prelimbic mPFC in mice with endogenous expression of Channelrhodopsin-2 (ChR2) in neurons expressing the vesicular GABA transporter (VGAT:ChR2). As optogenetic activation of interneurons has been shown to inactivate cortical network activity (Guo et al. 2014), this would allow for inhibition of mPFC during distinct task epochs (Fig. 3). We trained VGAT:ChR2 mice on both the AET (Fig. 3A) and a modified variable delay-to-signal task (VDST, Leite-Almeida et al. 2013), for comparison (Fig. 3B). In the AET, we silenced mPFC during the variable cue period (Fig. 3C) where we hypothesized attention is engaged most heavily, and observed a reduction in performance with continuous light stimulation (Fig. 3D) without a significant change in reaction time. To assess whether our reduction in performance could be reflecting an inability to report the cue in a matter unrelated to attentional load, we utilized the VDST, where we would predict attentional load should be greater in a different task epoch, the delay. When we photo-inhibited mPFC during the fixed cue period in the VDST (Fig. 3E), there were no differences in the percentage of correct choices or reaction time (Fig. 3F). However, when we silenced mPFC activity during the variable delay period (Fig. 3G), light stimulation reduced performance (Fig. 3H), without a significant difference in reaction time. This suggests that the AET is linked to the mPFC, and that mPFC activation is critical during cue detection in an attention-dependent manner.

**Figure 3.**
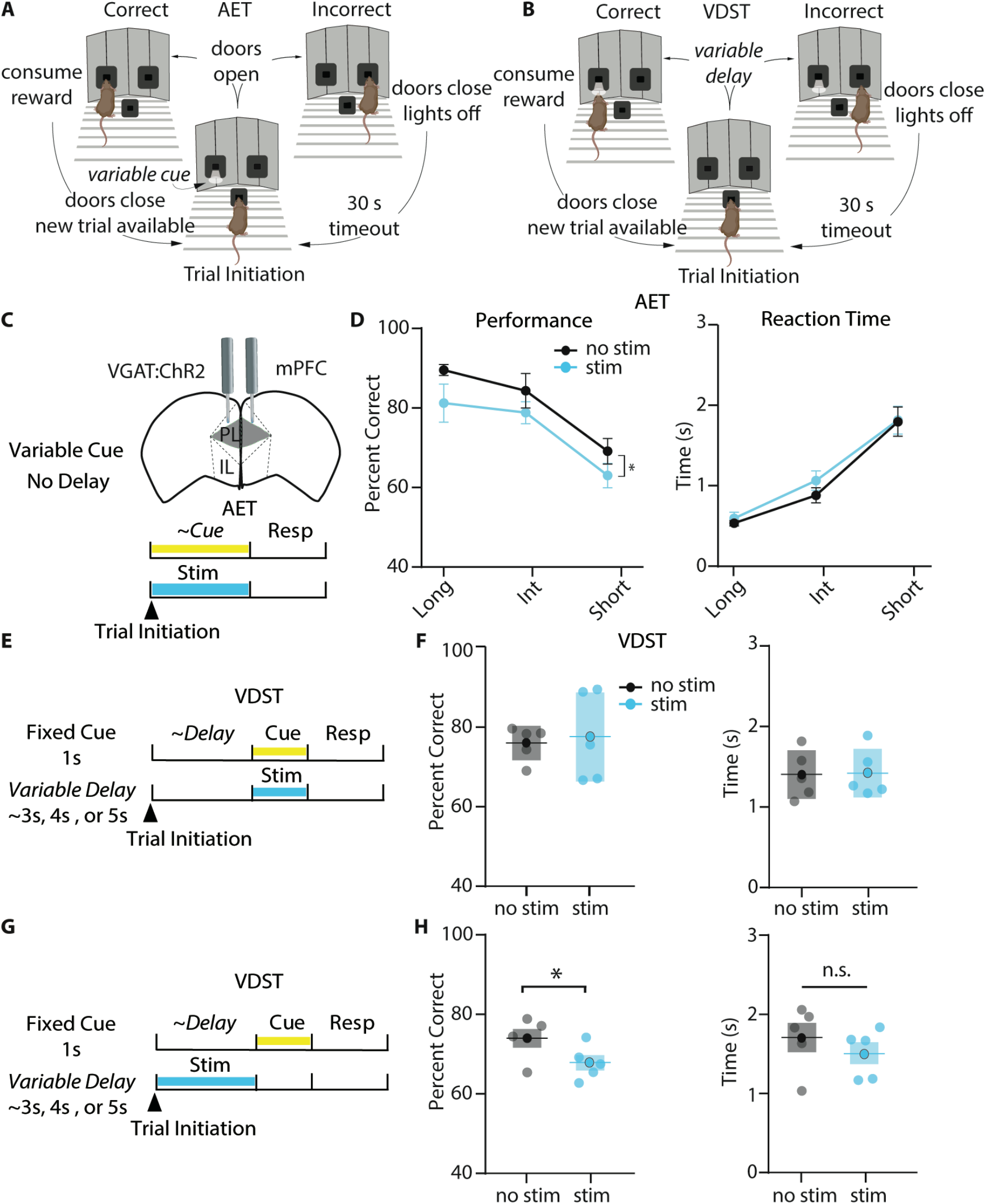
Cue detection in the AET is mPFC-dependent and linked to attentional load. A-B, Task design for the AET and VDST task. C, Experimental approach for disrupting prelimbic mPFC activity during the variable cue period with continuous blue light in VGAT:ChR2 mice. D, Photoinhibition with blue light reduces performance (left, two-way RM ANOVA, F_(1,18)_ = 6.990, p = 0.0165, n = 6), without affecting reaction time (right, two-way RM ANOVA, F_(1,18)_ = 3.323, p = 0.0850, n = 6) in the AET. E, Approach for photoinhibition during the fixed cue period in the VDST task. F, Performance or reaction time is not altered by light stimulation during the cue in the VDST (performance, paired t test, p = 0.7695; reaction time, paired t test, p = 0.6508; n = 6 for both). G, Approach for delivering continuous light stimulation during the variable delay period. H, Stimulation during the delay reduces performance in the VDST (paired t test, p = 0.0140) without altering reaction time (paired t test, p = 0.1268).

### Scn8a^+/−^ mice exhibit a loss of GABAergic inhibition in mPFC slice recordings

Our optogenetic experiments indicated the AET relies on mPFC function, suggesting a potential frontal locus for performance impairments in *Scn8a*^+/−^ mice. In patients, SWD is prominent over the frontal cortex, also suggesting frontal dysfunction (Archer et al. 2003, Tucker et al. 2007). Additionally, individuals with absence epilepsy exhibit reduced functional activation of the PFC that correlates with attentional impairments (Killory et al. 2011). Given this, we began with an *in vitro* approach to test for *Scn8a*^+/−^ related dysfunction in the prelimbic mPFC. We recorded network activity using evoked local field potentials (LFPs) in coronal slices collected from *Scn8a*^+/−^ mice and WT littermates (Fig. 4A, B) and derived a measure of current source density (CSD, Fig. 4C) (Freeman et al. 1975). This revealed several sinks or regions where cations flow into cells and out of the extracellular space, such as at excitatory synapses. We also observed nearby current sources, commonly attributed to the return flux of positive ions out of cells and back to the extracellular space. Of note, sources can also indicate sites of inhibitory synapses where anions flow into cells, with nearby sinks reflecting return paths. We developed a pharmacological approach to identify these distinct CSD components (Fig. 4D-H), and found a significant reduction specifically in one feature – a late source present over Layer 2/3 (Fig. 4H). Pharmacological characterization indicated that this source represents disynaptic GABAergic inhibition given its sensitivity to glutamatergic post-synaptic blockers, DNQX and CPP (Fig. 4-Supp. 1), and GABAergic modulator, Clonazepam (Fig. 4I), suggesting an attenuated recruitment of local mPFC GABAergic interneurons in *Scn8a*^+/−^ mice.

**Figure 4.**
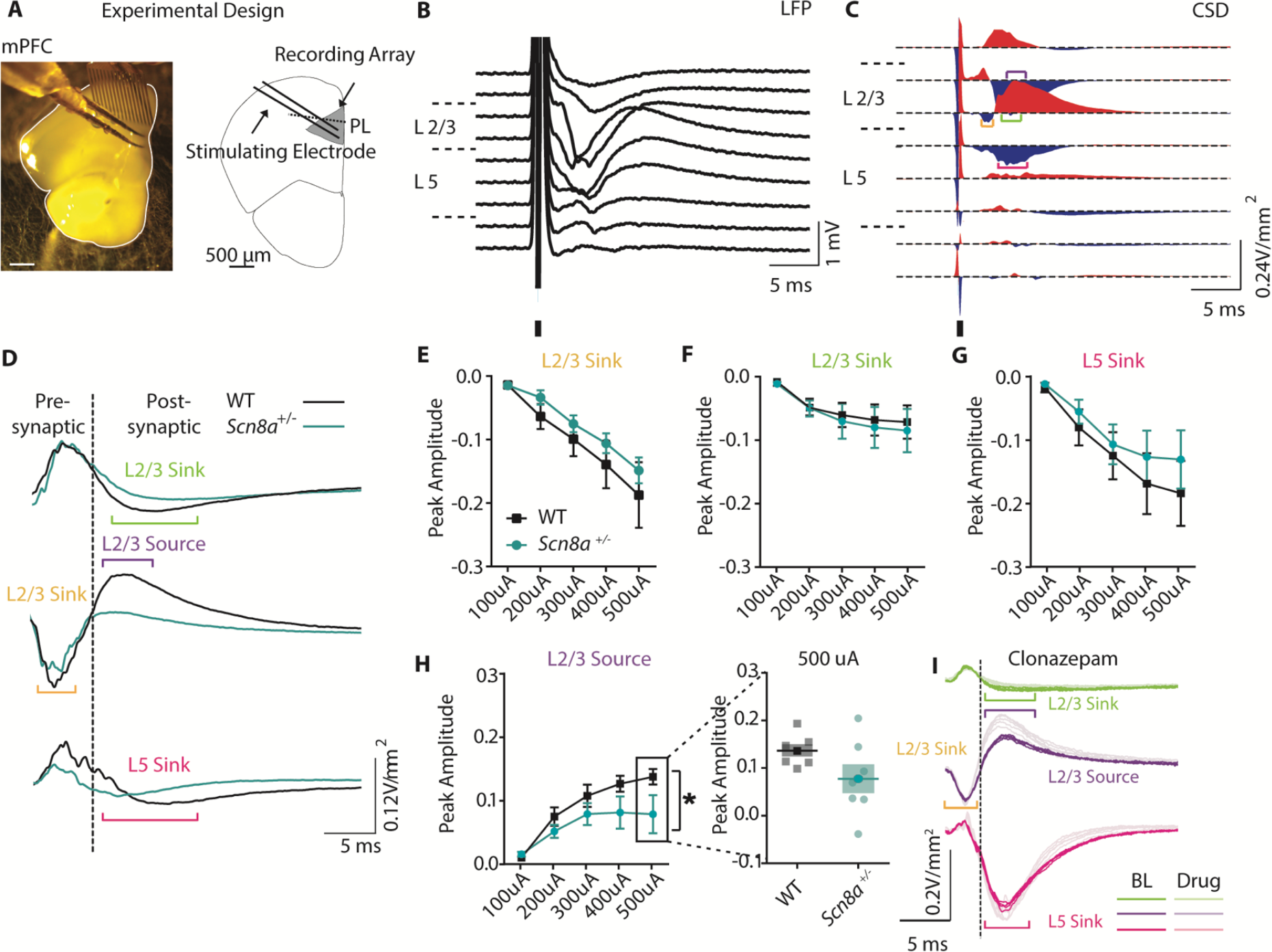
Scn8a^+/−^ exhibit a loss of GABAergic inhibition in mPFC slice recordings. A, Experimental design of LFP recordings. B, Representative local field potential (LFP) recording in response to a 250 μA electrical pulse. C, Representative current source density (CSD) derived from the LFP recording. Brackets highlight the traces used for quantification (orange = L2/3 sink, purple = L2/3 source, green = L2/3 sink, magenta = L5 sink). D, Representative responses of the pre- and post-synaptic L2/3 sink, L2/3 source, and L5 Sink to 250 μA electrical stim. E-H, Peak amplitude curves (V/mm^2^) of the response to electrical stim with increasing intensity. H, Only the L2/3 source showed a significant reduction in peak amplitude (two-way ANOVA, group effect, F _(1, 60)_ = 8.125, p = 0.0060, n = 7 slices for both WT and *Scn8a*^+/−^ mice). I, Example of clonazepam’s (100 nM) effects on the CSD components. Solid lines represent baseline CSD data, while transparent lines indicate CSD after wash on of clonazepam. See also Figure 4 Figure Supplement 1.

### Cue-evoked PVIN activity is correlated with AET performance and reduced in Scn8a^+/−^ mice

To determine the participation of GABAergic interneurons during behavior, we utilized fiber photometry to capture bulk Ca^2+^ signals. We chose to monitor PVINs because their activity has been classically linked to feedforward inhibition, a type of disynaptic inhibition important for gating cortical responses (Swadlow 2003, Delevich et al. 2015), as well as successful attentional performance and cognitively-associated gamma oscillations (Cardin et al. 2009, Sohal et al. 2009, Kim et al. 2016). We injected the mPFC of *Scn8a*^+/−^ mice and WT littermates homozygous for Cre expression in PVINs (*PV:Cre*) with an AAV5-CAG-FLEX-GCaMP6s virus and implanted an optical fiber for light delivery and collection (Fig. 5A). Following recovery from surgery, animals were trained on the AET as described above and GCaMP signals were collected throughout testing. The continuous signal (Fig. 5B) was segmented and aligned to trial initiation, and trials were sorted by cue length and accuracy (representative individual trials and average activity from WT and *Scn8a*^+/−^ mice, Fig. 5C,D). First, we took an average of each animal’s change in fluorescence (dF/F) during the baseline (the second immediately prior to the cue) and throughout the cue period for correct and incorrect trials and compared cue-related activity for all cue lengths. We found that for correct trials PVIN activity was consistently modulated at long and intermediate cue lengths (Fig. 5E), but not with the short, most challenging cue length. However, for incorrect trials, we found no significant modulation in PVIN GCaMP signals at any cue length.

**Figure 5.**
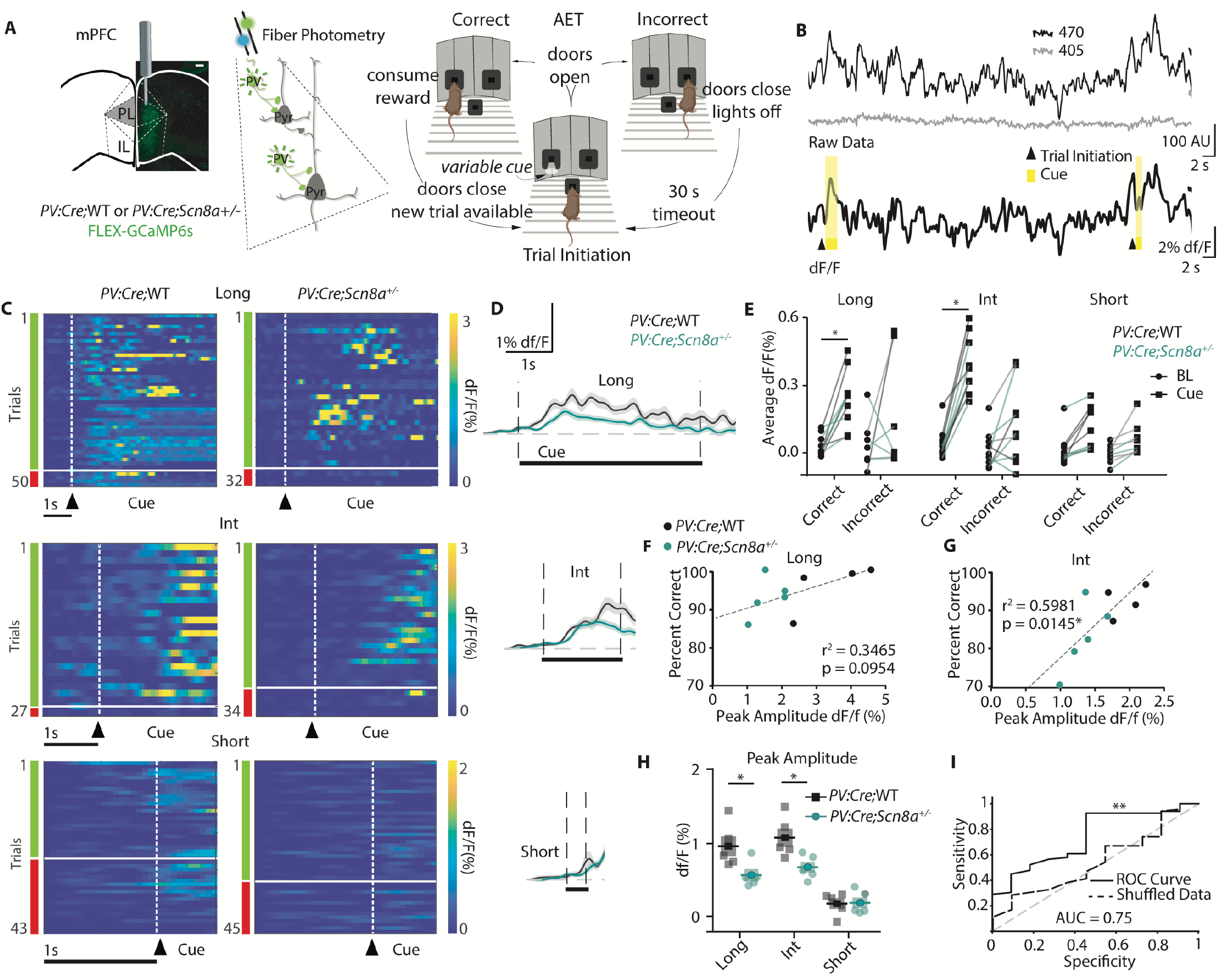
Cue-evoked PVIN activity is correlated with AET performance and reduced in Scn8a^+/−^ mice. A, *PV:Cre;* WT and *PV:Cre;Scn8a*^+/−^ were injected with a FLEX-GCaMP6s virus in the mPFC, and used for AET testing while recording Ca^2+^ signals from PVINs using fiber photometry. B, Top, example of the raw data collected from 470 (Ca^2+^-dependent) and 405 (Ca^2+^-independent isosbestic) excitation. Bottom, the same data after calculating the dF/F, with trial initiation and cue presentation indicated. C, Heat maps of all trials of dF/F activity and (D) average dF/F activity from representative animals in both groups and across cue lengths. E, Average dF/F baseline and cue activity at each cue length with groups indicated by line color. There was a significant interaction between the cue presentation and cue length in PVIN FiP signals (two-way RM ANOVA F _(4, 45)_ = 9.956, n = 10) during correct trials that was not observed in incorrect trials (F _(4, 45)_ = 1.680, n = 10). PVIN activity increased during the long and int cue in correct trials (p < 0.0001 for each), but not for the short cue or incorrect trials at any cue length. F, Data points represent each animal’s average performance at the long and int cue (G) plotted against the peak amplitude. All data points (*PV:Cre;WT* and *PV:Cre;Scn8a*^+/−^) were fit with a linear regression line. Peak amplitude was not a significant predictor of the percentage of correct choices for the long cue (r^2^ = 0.3465, p = 0.0954) but was for the int cue (r^2^ = 0.5981, p = 0.0145). H, Peak amplitude was reduced for *PV:Cre;Scn8a^+/−^* mice compared to *PV:Cre;*WT (two-way ANOVA, F _(1, 24)_ = 11.06, p = 0.0028, n = 5 for both *PV:Cre;*WT and *PV:Cre;Scn8a*^+/-^ mice) at the long (p = 0.0210) and int cue (p = 0.0210). I, A random forest classifier trained on the features from all FiP data for the int cue was able to predict trial outcome at a level significantly greater than chance as shown in the ROC-AUC curve (sensitivity vs specificity rate, p = 0.0056). See also Figure 5 Figure Supplement 1–3.

To determine what features of the cue-evoked signal were most important for accuracy we fit a linear regression model that attempted to explain average performance for all data at each cue length by either peak amplitude of dF/F, average dF/F, or the time to peak dF/F (Fig. 5-Supp. 1). Peak amplitude and time to peak were each significantly correlated with performance (Fig. 5-Supp. 1A and D), so these measures were utilized for group comparisons. For peak amplitude, only the intermediate cue peak amplitude was significantly correlated with accuracy (Fig. 5F, G and Fig. 5-Supp. 1A).

When we compared these features of cue-evoked PVIN activity during correct trials between groups, the peak amplitude was significantly reduced in *Scn8a*^+/−^ mice at long and intermediate cues, but not the short cue (Fig. 5H). Average dF/F was decreased in *Scn8a*^+/−^ mice at the intermediate cue (Fig. 5-Supp. 1C), however, the time to peak was not different between groups (Fig. 5-Supp. 1E). This suggests that generally increases in PVIN activity co-occur with accuracy (as it is observed with the long and intermediate cue where performance is higher), but only where attention is most heavily and variably recruited does it correlate with overall performance. To examine this specific reliance on PVIN activity at the attention-related intermediate cue more closely, we asked whether these measures could predict accuracy on a trial-by-trial basis. We trained a classifier using training data with the features peak amplitude and time to peak from all trials at each cue length, and evaluated its performance on a subset of held-out data. With long and short cues, our classifier was unable to predict trial outcome above chance (Fig. 5-Supp. 2). However, for the intermediate cue (Fig. 5I), the classifier correctly identified correct trials with 92% precision. We generated a receiver operating characteristic (ROC) graph, calculated the area under the curve (Fawcett 2006), and found that the AUC score was significantly greater than that generated by performing the identical classifier training procedure on randomly shuffled data. Taken together, this suggests that cue-evoked increases in PVIN activity may be critical for accurately responding to the intermediate cue and attentional engagement, as reductions in this activity are associated with incorrect trials and poorer performance in *Scn8a*^+/−^ mice. However, this relationship is distinct from the performance dependency observed for the long and short cues. Long cues evoke increases in PVIN activity that do not correlate with performance or lead to significant impairments in performance in *Scn8a*^+/−^ mice. Short cues which are associated with poor performance across genotypes cause no relative increase in PVIN activity compared to long or intermediate cues.

To explore whether decreases in PVIN activity were specific to cue perception, we also compared the peak amplitudes of reward-related activity between groups and observed that *Scn8a*^+/−^ mice exhibited reductions in PVIN activity during the time of reward consumption compared to WT mice (Fig. 5-Supp. 3A). However, reward activity was not correlated with average performance across groups (Fig. 5-Supp. 3B-D).

### Scn8a^+/−^ mice exhibit reductions in mPFC gamma power along with AET performance deficits

Our previous findings indicated reductions in cue-evoked PVIN activity in *Scn8a*^+/−^ mice. Given that analysis of ECoG/EEG remains the gold standard for diagnosis of absence epilepsy, we were curious if ECoG recordings might facilitate identification of network-level biomarkers related to attention dysfunction that could be measured in humans. We implanted a cohort of *Scn8a*^+/−^ and WT mice with a gold electrode over the mPFC and recorded ECoG (Fig. 6A) while mice performed the AET. As described above, we performed wavelet decomposition to obtain ECoG spectral power across different frequency bands (Fig. 6B) and compared power between *Scn8a*^+/−^ mice and WT littermates. AET performance was reduced in *Scn8a*^+/−^ mice (Fig. 6C) with concomitant reductions in spectral power for both the long and intermediate cue (Fig 6D,E). Specifically for the intermediate cue, there was a significant reduction in high gamma activity (60-90 Hz) in *Scn8a*^+/−^ mice. For the short cue we observed no differences in power across any frequency band (Fig. 6F), which is consistent with the absence of behavioral differences or evoked PVIN activity at this cue length.

**Figure 6.**
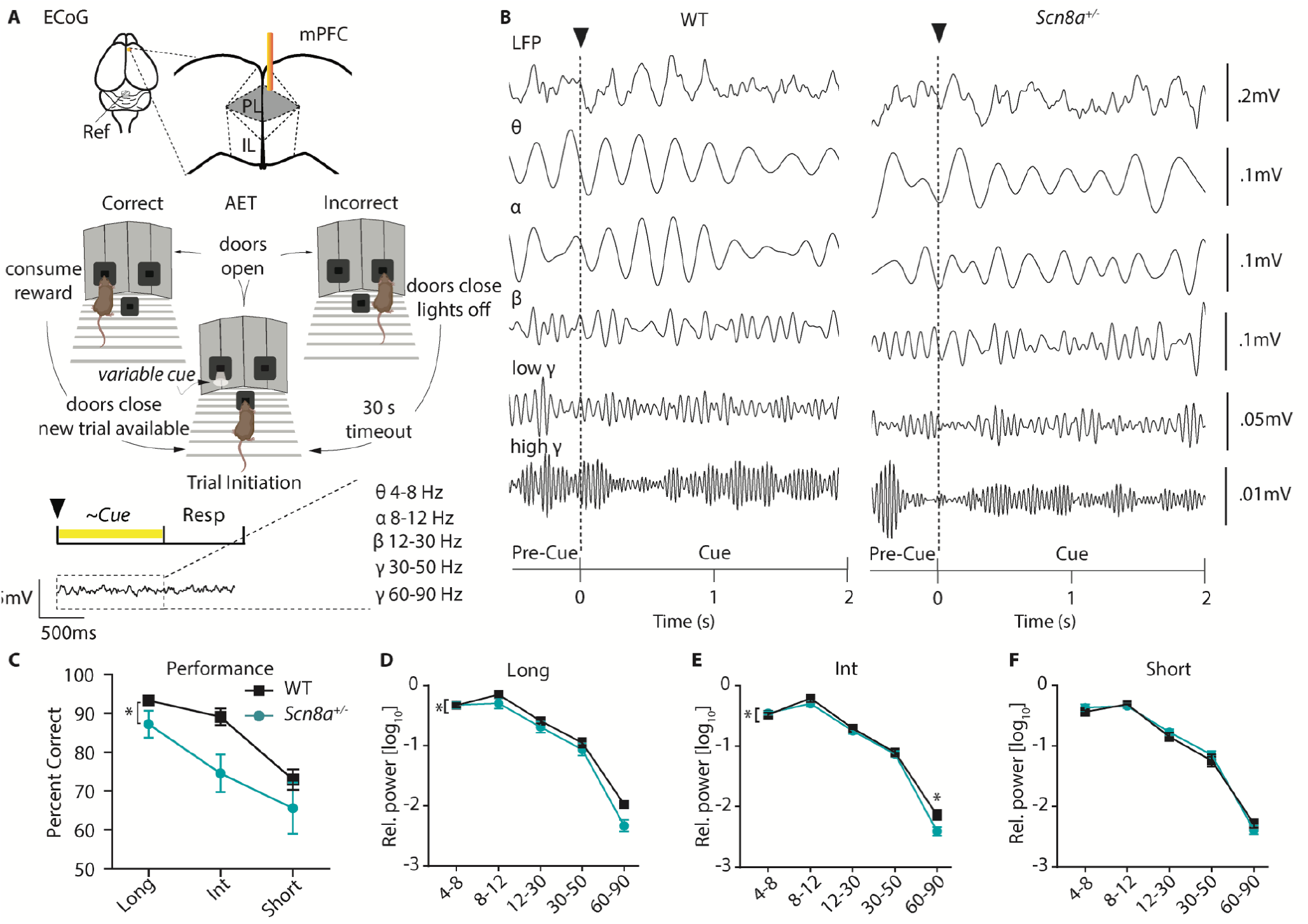
*Scn8a*^+/−^ mice exhibit reductions in mPFC gamma power along with performance deficits in the AET. A, Experimental design, ECoG activity was recorded using a gold electrode over mPFC and a reference screw over the cerebellum during the AET. Power was measured during the cue period across frequency bands. B, Representative ECoG filtered across different frequency bands during the cue period. C, Performance was decreased in *Scn8a*^+/−^ mice (two-way ANOVA, F _(1, 21)_ = 4.497, p = 0.0460, n = 3 and 6 for WT and *Scn8a*^+/−^ mice respectively). D-F, *Scn8a*^+/−^ mice have reduced power during the long (high gamma, p = 0.0597) and int cues (high gamma, p = 0.0145), but not during the short cue (long, two-way ANOVA, F _(1, 35)_ = 5.623, p = 0.0234; int, two-way ANOVA, F _(1, 35)_ = 4.586, p = 0.0393; short, two-way ANOVA, F _(1, 35)_ = 0.2656, p = 0.6095; n = 3 and 6 for WT and *Scn8a*^+/−^ mice respectively).

### Optogenetic activation of PVINs improves performance in Scn8a^+/−^ mice while common AED, VPA has no effect

PVINs are known to be critical in the generation of gamma oscillations (Cardin et al. 2009, Sohal et al. 2009) and activation of PVINs in the gamma range has produced beneficial effects on attention and cognition (Cho et al. 2015, Kim et al. 2016, Cho et al. 2020). Given this and the observed reduction in gamma power and performance at the intermediate cue in *Scn8a*^+/−^ mice, we hypothesized that supplementing gamma optogenetically may improve attention. Thus, we expressed channelrhodopsin2 (AAV5-FLOX-ChR2-mCherry) in mPFC PVINs of *PV:Cre;Scn8a*^+/−^ mice and implanted optical fibers for light delivery (Fig. 7A). Activation of PVINs at gamma frequency increased the percentage of correct choices for *PV:Cre;Scn8a*^+/−^ mice specifically at the intermediate cue (Fig. 7B). Importantly, this effect was not observed in *PV:Cre;Scn8a*^+/−^ mice that underwent the same experimental protocol with a control eYFP virus (Fig. 7-Supp. 1A).

**Figure 7.**
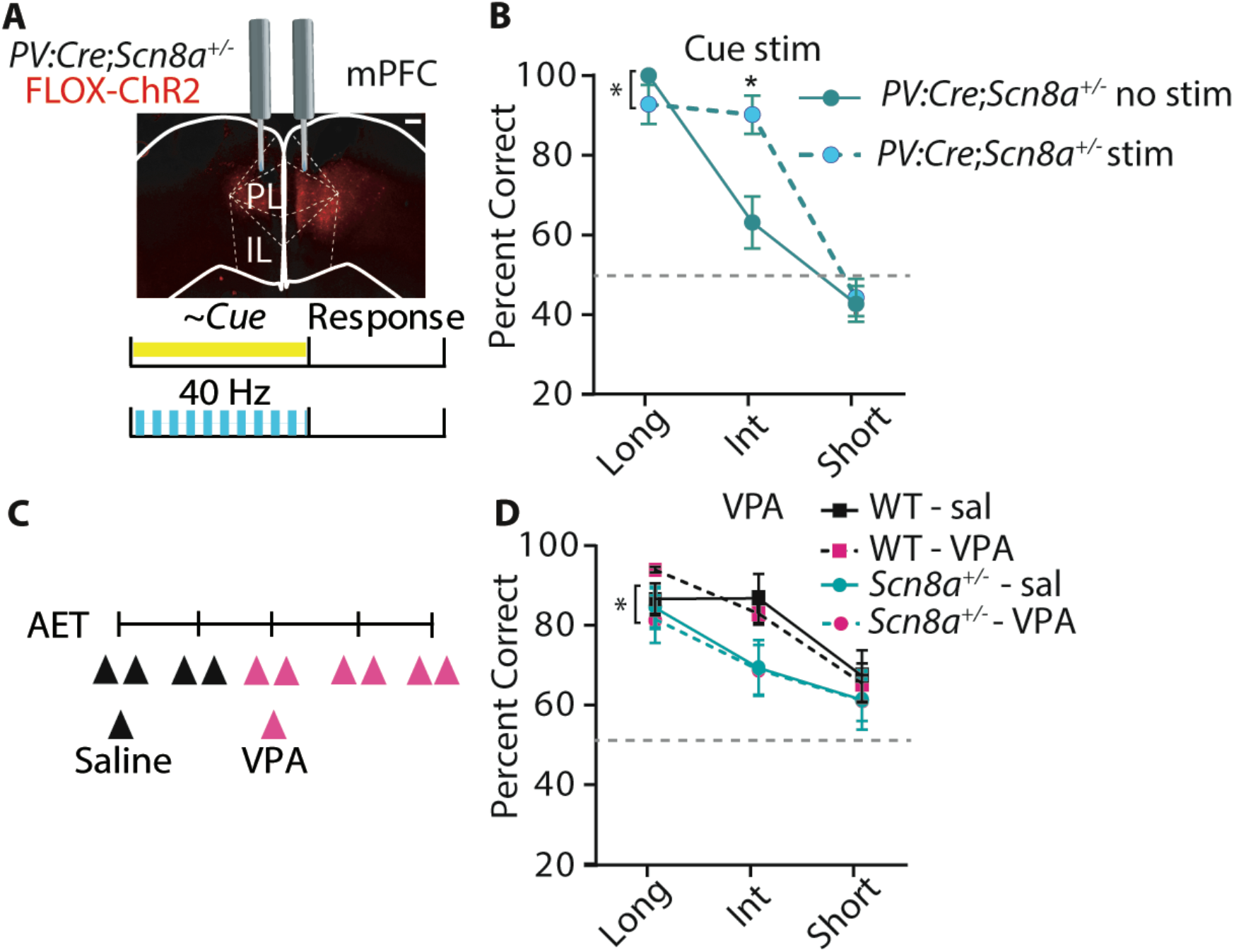
Optogenetic activation of PVINs improves performance in Scn8a^+/−^ mice while common AED, VPA has no effect. A, Mice received a bilateral injection with the FLOX-ChR2 virus, and then were implanted bilaterally with optical fibers for delivering blue light during the cue (40 Hz, 5ms pulse width). B, With gamma stimulation, there is a significant effect of light (two-way ANOVA, F _(1, 22)_ = 6.973, p = 0.0173, n = 7 mice) during the intermediate cue (p = .0044). C, Timeline of saline and VPA injections (200 mg/kg) during AET testing. D, While *Scn8a^+/−^ mice* had reduced performance in comparison to WT, there was no effect of VPA on performance (three-way ANOVA, group effect, F _(1, 35)_ = 5.850, p = 0.0191, VPA effect, F _(1, 35)_ = 0.010, p = 0.943; n = 3 and 6 for WT and *Scn8a*^+/−^ mice respectively). See also Figure 7 Figure Supplement 1.

We also were curious whether the commonly used anti-epileptic drug, Valproic Acid (VPA) would modify performance in the AET, given the lack of efficacy in mitigating cognitive symptoms in patients (Fig. 7C). WT and *Scn8a*^+/−^ mice were treated twice daily with saline and VPA, the latter of which significantly reduced seizure burden in *Scn8a*^+/−^ mice (Fig. 7-Supp. 1D). Although *Scn8a*^+/−^ mice exhibited reduced performance compared to WTs, there was no effect of VPA on accuracy in either group (Fig. 7D).

## Discussion

Our findings demonstrate that cue-evoked mPFC PVIN activity is strongly linked to and in certain cases, predictive of correct choices during a novel attentional engagement task (AET). Additionally, PVIN activity is positively correlated with performance both in WT and *Scn8a*^+/−^ mice. To our knowledge, this is the first demonstration of a cell type-specific activity pattern during rather than prior to cue perception predicting trial outcome during an attention task. These results also provide the first evidence of both the behavioral manifestation and a circuit biomarker of attention impairments in an animal model of absence epilepsy, one of the most commonly presenting cognitive comorbidities associated with the syndrome. Further, our findings support the recruitment of PVINs and gamma activity as a universal mechanism for visual discrimination and successful goal-directed behavior during attentional engagement.

### Attentional and cognitive impairments in absence epilepsy

Absence epilepsy is classically defined as an interruption of consciousness due to the emergence of stereotyped spike-and-wave activity ranging from 3-6 Hz in clinical populations (Watanabe 2003). However, patients also present with profound cognitive and behavioral impairments (Williams et al. 1996, Pavone et al. 2001, Caplan et al. 2008) with attention being one of the most severely impacted comorbidities (Fonseca Wald et al. 2019). These attention impairments are present even in interictal periods (Conant et al. 2010, Vega et al. 2010, D’Agati et al. 2012) and do not respond to current anti-epileptic drugs (Glauser et al. 2010, Masur et al. 2013). Gaining a better understanding of attentional dysfunction is critical for improving quality of life in patient populations, thus this study focused on investigating underlying mechanisms in an *Scn8a*^+/−^ mouse model that captures aspects of human generalized absence epilepsy (Berghuis et al. 2015).

Using our novel attention assay the AET, we confirmed that our model bore several similarities to the human condition. Most importantly we observed a consistent impairment in attention that appeared to exist independent of the seizure pathology. Seizures were reduced in *Scn8a*^+/−^ mice during task engagement, and treatment with VPA, a common AED, had no effect on AET performance. Along with observing no increase in power in seizure-related bands during incorrect trials in *Scn8a*^+/−^ mice or on average in comparison to WTs, we speculate that the attention impairments and seizures arise from distinct mechanisms.

### Medial PFC PVINs in attentional processing and attention dysfunction

One such mechanism may be the recruitment of mPFC PVINs. The PFC has long been implicated in attention across species. However, only with recent advances has it been possible to begin to answer questions about the contributions of genetically distinct cell-types to attentional performance. Optogenetic (Kim et al. 2016) and pharmacogenetic (Ferguson et al. 2018) manipulation of PVINs improves vigilance and attentional flexibility in rodents, raising the question of how PVINs support basic attentional processes. Using a task designed specifically to capture attentional engagement, we observed that by several measures population rises in PVIN activity were strongly associated with overall accuracy.

Cue-evoked PVIN activity was significantly reduced in *Scn8a*^+/−^ mice compared to that of WTs, and this was predictive of impairments in AET performance. This was corroborated by network electrophysiology experiments both *in vitro* and *in vivo*, providing a novel circuit-level mechanism for attention impairments in absence epilepsy. Reductions in feed-forward inhibition *in vitro* were found concomitant with a loss of gamma power during the AET, arguing for PVIN-mediated pathology in *Scn8a*^+/−^ mice. Importantly, correlated hypoactivity of PVINs and AET performance impairments as well as AET improvement with optogenetic stimulation in *Scn8a*^+/−^ mice were all observed at the intermediate cue length. This suggests that response variability at the intermediate cue specifically involves attentional processes and can be used to further elucidate cell-types and circuits supporting attention along with attentional dysfunction.

Questions remain about the source of the recruitment of PVIN activity, contributions of other inhibitory cell types to task components in the AET, and how top-down activity from the mPFC modulates activity in other structures to drive the appropriate behavioral output. Studies point to the involvement of the mediodorsal thalamus (Freeman et al. 1975, Delevich et al. 2015, Schmitt et al. 2017, Rikhye et al. 2018) in modulation of mPFC PVIN activity related to attention, so future studies will examine whether deficits in PVIN activity in absence epilepsy arise locally or are downstream of thalamic hypofunction. Of note, previous work in *Scn8a*^+/−^ mice suggests that relay cell function of another thalamic nucleus, the ventrobasal thalamus remains intact (Makinson et al. 2017). Regardless of their mechanisms of regulation or recruitment, broadly elevated mPFC PVIN activity seems to provide an optimal cortical state for improved information processing during attention and other higher-order cognitive behaviors. This may be due to their ability to suppress neurons representing distracting information (Ferguson et al. 2018) or their importance in the generation and maintenance of gamma oscillations (Cardin et al. 2009, Sohal et al. 2009). In support of this, gamma power was reduced in *Scn8a*^+/−^ mice, and gamma frequency stimulation of PVINs improves their attention.

### Beyond cognitive deficits in patients with absence epilepsy and animal models

Somewhat surprisingly, our behavioral tests revealed that other behaviors were essentially unaffected. While attention impairments present as one of the most consistent comorbidities, reductions in sociability and affective behaviors have previously been reported in patients (Rantanen et al. 2012) and rodent models, including the Wistar Albino Glaxo from Rijswijk (WAG/Rij) and Genetic Absence Epilepsy Rat from Strasbourg (GAERS) rat models (Sarkisova et al. 2011, Henbid et al. 2017) and mice with *Scn8a* mutations (McKinney et al. 2008, Papale et al. 2010). Reports from animal models suggest that ethosuximide, another commonly prescribed AED in absence epilepsy, may mitigate affective symptoms in models of absence seizures (Dezsi et al. 2013) and chronic pain (Kerckhove et al. 2019). Given that AEDs can yield improvement across some symptom domains (seizures and affective symptoms), this points toward a likely distinct mechanism of attention and cognitive impairments, as they remained resistant to VPA treatment in *Scn8a*^+/−^ mice. We did observe decreases in reward-related PVIN activity in *Scn8a*^+/−^ mice, and although this was not related to task performance, it could suggest more global PVIN hypofunction that may mediate other behavioral impairments in absence epilepsy. Additionally, *Scn8a*^+/−^ mice exhibited increases in repetitive responding during the AET which may indicate higher levels of behavioral inflexibility. Behavioral flexibility can present in various ways with distinct circuit contributions (Hamilton et al. 2015), thus future studies will examine the impact of *Scn8a* loss on set-shifting and reversal learning along with the potential involvement of the orbital frontal cortex, thalamus, and ventral striatum (Floresco et al. 2009). Additional studies are also needed to characterize the generalizability of how blunted PVIN activity contributes to other behavioral impairments in *Scn8a*^+/−^ mice and across rodent models.

### Implications for cognitive therapies in absence epilepsy and other disorders

As attentional engagement precludes information perception related to other cognitive domains, such as working memory, attentional shifting, and even information encoding for short or long-term memory, deficits in attention are likely to occur in concert with a host of cognitive abnormalities. In addition to absence epilepsy, attention impairments present as a comorbidity in diseases with complex cognitive behavioral phenotypes, including schizophrenia (Green 1996), autism (Allen et al. 2001), depression (Rock et al. 2014), bipolar disorder (Cullen et al. 2016), Alzheimer’s disease (Perry et al. 1999), and epilepsy (Holmes 2015). Of note, *Scn8a* mutations are also present in other disorders with complex cognitive phenotypes including epileptic encephalopathies, intellectual disability, and autism spectrum disorders (Larsen et al. 2015, Butler et al. 2017, Wagnon et al. 2017). If PVIN hypofunction continues to be linked the etiology in attention and cognitive dysfunction across disease states, moving forward, it will be critical to identify strategies for pharmacological or other avenues for cell-type specific augmentation of PVIN activity and/or gamma oscillations. This will hopefully accelerate development of better therapies for attention that could improve functional outcome in absence epilepsy and myriad neurological and psychiatric diseases.

## Abbreviations

mPFC: medial Prefrontal Cortex
PVIN: parvalbumin interneurons
AET: attentional engagement task
ECoG: electrocorticogram
CSD: current source density
ROC-AUC: receiver operating characteristic -area under the curve
GABA: gamma-amino-butyric acid
IR: infrared

## Author Contributions

BRF and JRH designed the research, BRF performed the experiments and analyzed the data, and BRF and JRH wrote the article. JRH supervised all aspects of the project.

## Materials and Methods

### Lead Contact

Further information and requests for resources and reagents should be directed to and will be fulfilled by the lead contact, John Huguenard.

### Materials Availability

This study did not generate any unique reagents.

### Data and Code Availability

Source data to recreate figures presented in this manuscript has been uploaded to Dryad and can be found here: https://dataverse.harvard.edu/dataset.xhtml?persistentId=doi:10.7910/DVN/MCMHSW. Custom MATLAB scripts used for analysis will be uploaded to github upon acceptance of the manuscript.

### Experimental Model and Subject Details

Mice with the heterozygous loss of function mutation in *Scn8a* were purchased from the Jackson laboratory (C3HeB/FeJ-Scn8amed/J, Kohrman et al., 1996, Stock#: 003798), referred to in this manuscript as *Scn8a+/−*. For the behavioral experiments, including the attentional engagement task (AET), social interaction, novel object recognition, and the anxiety task, *Scn8a*^+/−^ mice were crossed with C57BL/6J mice to result in litters that were either *Scn8a*^+/−^ or WT (*Scn8a-/-*). For fiber photometry and optogenetics experiments, *Scn8a*^+/−^ mice were crossed with mice homozygous for Cre expression in parvalbumin interneurons (B6;129P2-Pvalbtm1(cre)Arbr/J, *PV:cre,* Stock#: 008069) to generate *Scn8a*^+/−^ and *Scn8a-/-* mice, which were both heterozygous for *PV:cre*. For photoinhibition experiments, hemizygous VGAT-ChR2 mice were purchased from Jackson Laboratory (VGAT-mhChR2-YFP, JAX stock#: 014548) and bred with C57BL/6J mice to maintain the colony (Stock#: 000664). Only ChR2 positive mice were used for the photo-inhibition experiment. Mice of the desired genotype were randomly assigned to experimental groups. All mice were maintained on a reverse 12 hour dark/light cycle, and all experiments occurred during their active cycle. Males and females were used all experiments and were 2-4 months of age. No experimental differences due to gender were observed. Food and water were available ad libitum, except for during training and testing for the AET. For the AET and variable delay-to-signal task (VDST), mice were food restricted to approximately 85% of their free-feeding weight, two weeks prior to beginning experiments. All experiments were approved by the Stanford Administrative Panel on Laboratory Animal Care (APLAC, Protocol #12363) and were performed in accordance with the National Institute of Health guidelines.

### Method Details

#### Surgery

Mice aged postnatal day 40-50 were used for all surgeries. For the fiber photometry experiments, *Scn8a*^+/−^ or WT mice received a unilateral injection of a AAV5-CAG-FLEX-GCaMP6s (Addgene, #100842-AAV5) tagged with GFP into the prelimbic medial prefrontal cortex (mPFC) using the coordinates (AP: −2.6, ML: +0.25, DV: −1.0). Briefly, a Hamilton syringe was slowly lowered to the target region, and the virus (300 nL) was infused at a rate of 50 nL/min. The cannula remained in place for 5 minutes post injection, then an optical fiber (400 μm, 0.48-NA, 1.25 mm-diameter stainless steel ferrule, Doric Lenses) was implanted slowly directly above the viral infusion site. For optogenetics, the virus used was an AAV5-FLOX-ChR2-mCherry (Addgene, #20297-AAV5) or a AAV5-Ef1a-DIO-eYFP (UNC Vector Core), and the viral injection and optical fiber (200 μm, 0.22-NA, 1.25 mm-diameter stainless steel ferrule, Doric Lenses) insertion procedure was the same, except it was bilateral rather than unilateral. VGAT:ChR2 mice also were implanted with bilateral optical fibers into the mPFC. Following optical fiber insertion, the optical fiber was stabilized, and the incision was closed using dental cement. After surgery, animals recovered for one week before food restriction, and a week later, experiments began. Following experiment completion, animals were sacrificed for histological verification of viral expression and optical fiber placement. Animals with insufficient viral expression or improper optical fiber placement were excluded from analysis.

#### Behavior

Attentional Engagement Task (AET). For the AET, mice were food restricted for at least one week prior to experiments to increase motivation for the task. First, mice were habituated to the chamber, that was modeled from (Wimmer et al. 2015) and consisted of a center initiation port, and two reward ports on the left and right side. During training, mice learned to use visual cues (a white LED positioned below each reward port) to locate a food reward. During each session, mice remained in the chamber until 80 minutes had passed, or they had completed 100 trials, whichever came first. To initiate trials, mice learned to place their nose in a center portal, breaking an infrared light-emitting diode (IR LED) beam between an IR LED-photodiode pair. Following the visual cue (5s LED flash), doors blocking access the two reward ports would raise. A correct choice would trigger reward delivery (10 μl of condensed milk) from a syringe pump (World Precision Instruments) connected to the reward port. Following an incorrect choice, the doors would close, leading to a 30 second timeout, where the animal could not initiate another trial. All mice had to reach a criterion of at least 70 % correct for 3 consecutive training days, or 90% correct for 2 days. During testing, after trial initiation, a visual cue of either 5s, 2s, 1s, 500ms, and 100 ms, or 5s, 2s, or 500 ms, indicated the correct location of the food reward. The cue length and correct side varied pseudorandomly, and performance was averaged over 3 test days. Omissions were quantified as trials in which the animal took longer than 5s following cue termination to make a choice. Reaction Time was the time between trial initiation and making a response in one of the reward ports. In single-cue length blocks, trials were organized on to 10-trial blocks of each cue length from longest to shortest that repeated until the end of the session. Repetitive responding was quantified by counting the number of blocks of 5 consecutive trials in which the animal selected the same port normalized by the total trial number in a session. For the VGAT:ChR2 experiments, training for the VDST began immediately following the final AET test day. Animals were trained using the same strategy of reinforcement and punishment as described above to respond to fixed 1s cue following a 2s delay. To avoid overtraining, mice had to complete only one day at 60% correct to continue to testing. During testing, the delay prior to the cue varied between 3s, 4s, and 5s while the cue length remained fixed at 1. Trial logic was controlled by a microcontroller (Arduino Mega 2650), and custom MATLAB scripts were used to analyze data and determine the percentage correct at each cue length.

Sociability. To measure sociability, mice were tested in a three-chamber social interaction task (Moy et al., 2004). For the three-chamber paradigm, the apparatus was partitioned into three zones, with a small walkway so the mouse could move freely between chambers. A camera was mounted directly above the apparatus to capture video that was used for later analysis. Animals were habituated to the entire apparatus for ten minutes. Then, an empty cage was positioned in one chamber, a cage containing a novel mouse was placed in the opposite chamber, and the mouse was given ten minutes for exploration. Videos were scored by the experimenter, and active interaction, time spent oriented toward, sniffing, or interacting with the empty cage versus the cage with the novel mouse was measured.

Object Recognition. Object recognition was tested as previously described (Barker et al., 2007) in a clear plastic chamber (14 × 16 × 6”) in which mice were habituated to for 5 minutes on the day prior to beginning testing. One object pair was a set of four stacked bottle caps, alternating between purple and orange caps. The other was a glass jar of equal height as the tower of caps, filled with purple sand. The testing day consisted of a sample phase and a test phase, with a two-hour delay between. During the sample phase, mice were given 3 minutes to explore two identical objects placed in the center of the arena. For the test phase, one of the objects was replaced, and the animal was allowed to explore both objects freely for five minutes, while video was acquired from above the testing arena. The experimenter scored the videos by measuring the total time spent exploring each object within the first 20 seconds of total exploration. A difference score was calculated by taking the time exploring the object in phase 2 from phase 1, then dividing that difference by the total time (20 seconds). Heatmaps of the animal’s activity in the during object recognition and social interaction using ezTrack in Jupyter Notebooks (Pennington et al. 2021).

### Electrocorticography Recordings and Analysis

Mice were surgically implanted with bilateral ECoG screw electrodes into the skull above the somatosensory cortex (S1) or gold pin electrodes over the mPFC, and recorded channels were referenced to an ECoG screw electrode above the cerebellum. Data were acquired using the OpenEphys data acquisition board and software (Siegle et al. 2017). Data were bandpass filtered using a Butterworth filter (0.1 - 100 Hz) and sampled at 30 kHz. We recorded ECoG from animals over 3 sessions where they were initiating trials at a rate of at least 1 trial per minute (engaged in task). For the S1 recordings, this was compared to 3 one-hour home cage sessions. All data processing and analysis were performed using custom MATLAB scripts. Continuous wavelet transformation was used for spectral analysis (Torrence et al. 1998) as described in (Sorokin et al. 2017). For seizure detection, we utilized the full power spectrogram to detect seizures and events were only included if they were between 2 and 30 s in duration. Additionally, events that exhibited maximum power beyond the typical absence seizure spectrums were discarded. Seizures were normalized to session length, and displayed as seizures/minute. For power analysis, data was segregated into by cue length and accuracy, and mean power in the modified theta (Ø∗, 7-10 Hz) or β (10-20Hz) range which encompass the fundamental and 2^nd^ harmonic frequency bands for absence seizures (Sorokin et al. 2017) was extracted, and displayed as mean power and variance in relation to the cue. For the mPFC recordings, we extracted data from the cue period across trials and separated data by cue length and accuracy. We generated ECoG power spectrograms (1-100 Hz) using the continuous wavelet transform (Morlet wavelet in the Matlab wavelet toolbox). The spectrograms were normalized to peak, and power was averaged within bands of interest (4-8, 8-12, 12-30, 30-50, 60-90).

### Extracellular Multi-Unit Electrophysiology

To measure differences in network activity, extracellular recordings were obtained from mice who had been previously used for fiber photometry experiments. Mice were anesthetized with pentobarbital (50mg/kg) and perfused with ice-cold sucrose buffer containing (in mM): 234 sucrose, 2.5KCl, 1.25 NaH2PO4, 10 MgSO4, 0.5 CaCl2, 26 NaHCO3, and 11 glucose, equilibrated with 95% O2 and 5% CO2, pH 7.4). Mice were decapitated, the brain was extracted, and coronal slices (400 μm) were collected using a Leica VT1200S vibratome. Slices were transferred to oxygenated ACSF (containing in mM: 126 NaCl, 2.5 KCl, 1.25 NaH2PO4, 2 MgCl2,2 CaCl2, 26 NaHCO3), that bubbled continuously with 95% O2 and 5% CO2, and incubated at 32°C for 1 hour. Then, slices were incubated at room temperature for 1-5 hours before they were placed in an interface recording chamber, in which they were continuously perfused with oxygenated ACSF (30-32°C) at a flow rate of 2-3ml/min. A linear silicon multichannel probe (16 channels, 100 μm inter-electrode spacing, NeuroNexus Technologies) was placed in the prelimbic mPFC perpendicular to the laminar plane, such that the electrode array collected local field potentials (LFPs) from each cortical lamina. A bipolar tungsten electrode was placed immediately ventral to the recording array in layer II/III to deliver electrical pulses (0.1 ms, 100-500 μA in 100 μA increments) to evoke synaptic responses. Signals from all sixteen channels were digitized at 25 kHz, using a 3000 Hz lowpass filter, amplified and stored using a RZ5D processor multichannel workstation (Tucker-Davis Technologies). To better examine the location, direction, and magnitude of currents evoked in response to electrical stimulation, a current source density (CSD) analysis was performed by calculating the second spatial derivative of the LFP (Freeman et al. 1975). When net positive current enters a cell, this creates an extracellular negativity that is reflected in a current “sink,” and appears as a negative deflection in the CSD. Conversely, current “sources” indicate net negative current flowing into a cell and will create positive CSD responses. Differentiation of CSD components into pre and post-synaptic as well as determining distinct neurotransmitter contributions were identified. DNQX (25 μM, Sigma, D0540) and CPP (1 μM, Sigma, C104) were used to determine disynaptic and post-synaptic components, while Clonazepam (a benzodiazepine, 100 nM) was used to identify components reflecting or related to GABAergic neurotransmission. Peak amplitudes were quantified using trial-averaged responses at each stimulation intensity (10 trials each). LFP and CSD plots were generated using custom MATLAB scripts.

### Fiber Photometry

Fiber Photometry was used to measure aggregate Ca^2+^ signals from GCaMP6s-expressing PVINs during the AET. The set up was designed as described in (C. K. Kim et al., 2016). Briefly, an implanted optical fiber (1.25 diameter stainless steel ferrule) was coupled to a patchcord with ceramic sleeves (Thorlabs, ADAL1). The patchcord terminated in an SMA connector (Thorlabs, SM1SMA), which was focused onto the sensor of an sCMOS camera (Hamamatsu, ORCA-Flash4.0). A series of emission filters and dichroic mirrors were arranged such that 470 or 405 nm light from two LEDs could be transmitted through the patchcord, and 535 nm light would be collected at the working distance of the objective. A custom MATLAB GUI (https://github.com/deisseroth-lab/multifiber) triggered the LED drivers (Thorlabs, LEDD1B) through a National Instruments Data Acquisition device (NIDA, PCIe-6343-X). This GUI triggered alternating pulses of the GCaMP activating or control wavelength (470 and 405 respectively) at a frequency of 40 Hz to result in collection of Ca2+ and isosbestic signals at a 20 Hz sampling rate. To account for motion-artifacts, the reference trace (405 nm) was scaled to the GCaMP (470 nm) using a least-squares regression, then we subtracted the scaled reference from the GCaMP signal to get a normalized trace. To obtain the change in fluorescence over time (dF/F), we subtracted the median of the normalized 470-nm signal from the continuous signal at each point in time, divided by the median ([f(continuous) - f(median)/f(median)). To align the dF/F to behavior, the Arduino sent TTL pulses at the start of each trial to the NIDA device for storage in a MATLAB file along with the fiber photometry data. The dF/F was separated into trials by cue length and accuracy, and peak, average, and time-to-peak were quantified using custom MATLAB scripts. Representative heatmaps of individual trial data were generated using the MATLAB function imagesc.

### Machine Learning Trial Prediction

Trial Prediction Analysis was performed in python using built-in functions from the *sklearn* and *imblearn* toolkits. Data were organized into tables with trials for each cue length with the features peak and time-to-peak for each trial along with its classifier, correct or incorrect. Data were separated into training and testing data sets using an 80/20 split. Given that there were often much more correct than incorrect trials at longer cue lengths, the incorrect trials (minority class) were oversampled using the synthetic minority oversampling technique (Chawla et al. 2002), then the features from the training and test set were normalized independently. The performance of several classifiers was evaluated on the training data using repeated K-fold cross-validation (k = 10, 10 repeats) to get the average f1-scores and receiver operating characteristic-area under the curve (roc-auc) scores. Based on this, a random-forest classifier was chosen to make predictions on the test set. To test classifier skill, we generated ROC-AUC curves on the test set, and compared that with curves generated using the identical training and testing procedure described above but with random shuffled data.

### Optogenetics

To deliver optogenetic stimulation during behavior, we utilized several components of the fiber photometry system described above including the LED and LED driver, patchcord, and optical fibers. Once the animal initiated a trial, the Arduino sent TTL pulses directly to the LED driver, such that blue light pulses (470 nm, 40 Hz, 5ms pulse width, 0.5mW) were delivered to through the patchcord to the mPFC throughout the duration of the cue.

### Pharmacology

For evaluation of VPA treatment on AET performance and seizures, animals were treated twice daily with the following schedule: Twice daily saline beginning in the evening two days before testing, and continuing for two days of testing, then twice a day VPA (200 mg/kg) beginning two evenings before day one of VPA testing, and continuing for three days. ECoG was recorded during each behavioral session. Data was analyzed from two days of saline testing and three days of VPA testing and averaged for comparison. Animals were removed from food restriction and allowed a two-week wash out period, then this same approach was repeated without behavioral testing. Animals were placed in the behavioral chamber for one-hour ECoG sessions while satiated. Seizures were detected as described above in each session, averaged over sessions (two saline and three VPA), and displayed as seizures/minute in saline versus VPA.

### Histology

All animals with exception of those for electrophysiological recordings were sacrificed for histological confirmation of viral injection and expression. Mice were transcardially perfused with 0.1 M PBS followed by 4% paraformaldehyde (PFA) in PBS. Brains were removed and postfixed for 48 hours, then cryoprotected in 10% to 30% sucrose solution for an additional 48 hours. Forty μm sections were collected using a vibratome, and sections were mounted and cover-slipped with VECTASHIELD ® Antifade Mounting Medium with DAPI (Vector Laboratories, Burlingame, CA). Images were acquired with a ZEISS ApoTome 2.

### Quantification and Statistical Analysis

All statistical analysis was performed using MATLAB, Prism 6 (GraphPad), and python using the Spyder IDE. We determined sample sizes using a power analysis along with our previous studies and literature. All measures are presented as mean ± SE or individual data points. For all behavior experiments, n represents individual animals. In slice experiments, n represents number of slices with no more than 2 slices per animal. For fiber photometry, electocorticography, and optogenetics experiments, n represents AET data averaged over 3 days from individual animals. Normality was determined using the Shapiro-Wilk test to direct the usage of parametric versus non-parametric tests. Outliers were determined using the mean absolute deviation method and any data set where outliers were identified is noted in the source data files. For normal data, the comparisons of two independent groups were done using a Student’s t test, while paired samples were compared using paired t tests. For data without a normal distribution, a Mann-Whitney U test (independent) or Wilcoxon Signed Rank test (paired) was used. For comparisons of multiple groups and cue lengths, a two-way ANOVA was used followed by post-hoc Holm-Sidak’s test with correction for multiple comparisons. P < 0.05 was considered significant and was represented by a one asterisk (“*”), while p < 0.01 was represented with two (“**”). The experimenter was blinded to group identity during data all data collection and analysis.

## Supplemental Figures and Legends

**Figure 2 Figure Supplement 1.**
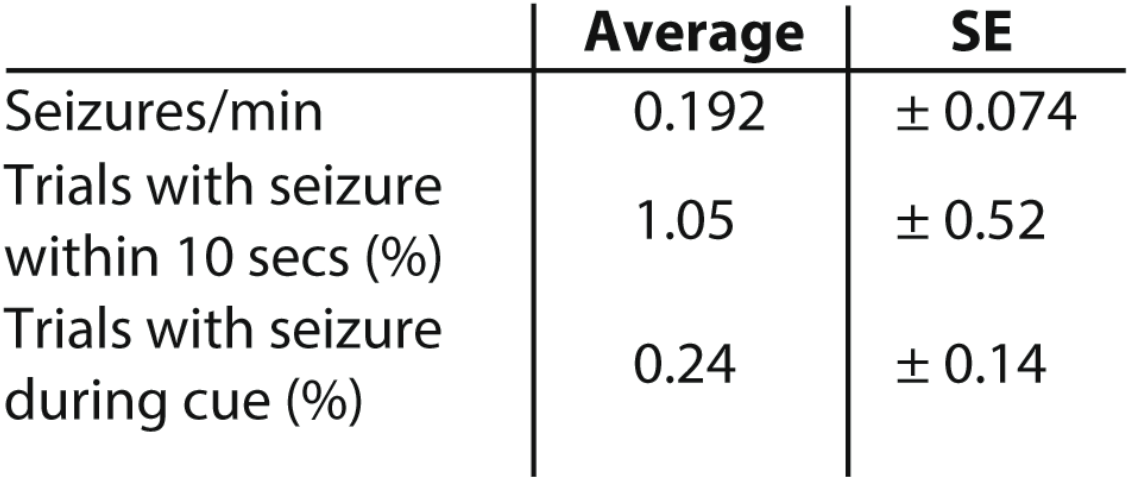
Quantification of seizure frequency during the AET. Seizures/min (example seizure in figure 2) during the task, percent of trials with seizures during cue, and percent of trials with seizure within 10 seconds of trial are presented. All measurements represent average totals for animals across three behavioral sessions. N = 6 for all measurements.

**Figure 4 Figure Supplement 1.**
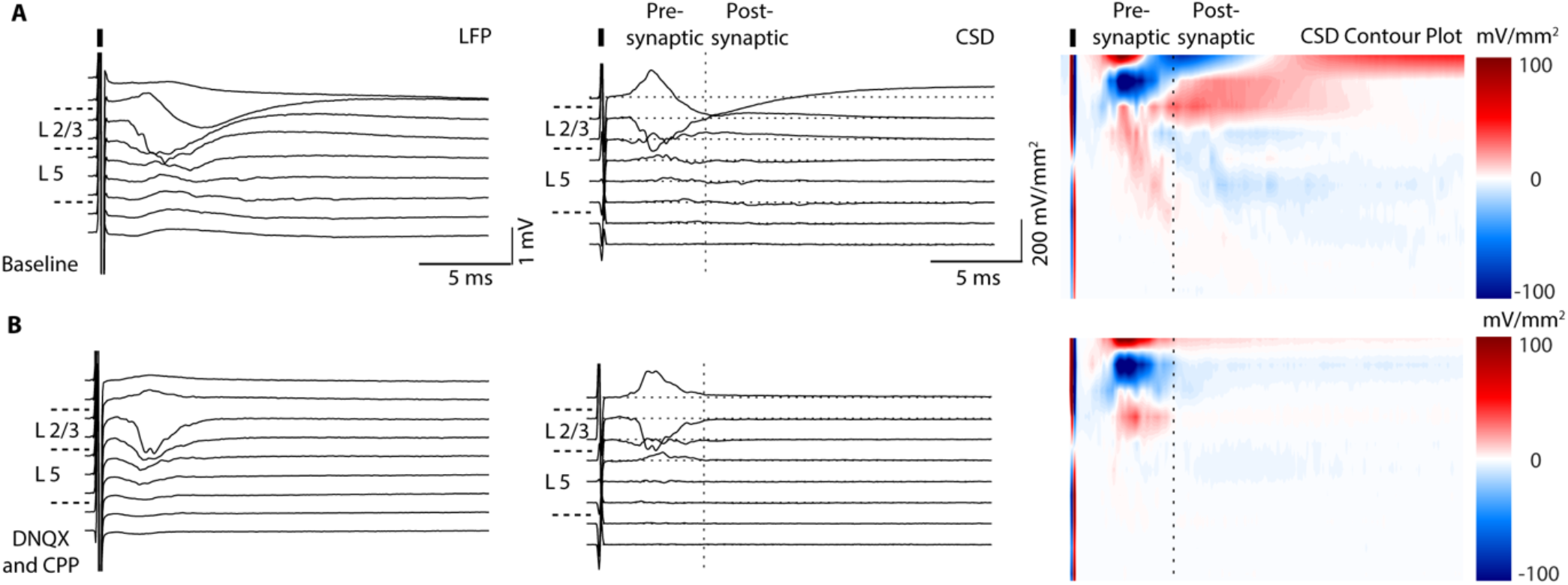
DNQX and CPP reduce late activity in the mPFC LFP recordings. A, Left, Representative baseline local field potential (LFP) recording in response to a .1 ms 250 μA electrical pulse, CSD (middle), and contour plot (right, sinks in blue, and sources in red). B, Pre-vs. post-synaptic activity were determined by the reduction of the late CSD components by DNQX (25 μM) and CPP (1 μM). Data are shown as averages over 10 trials in each condition.

**Figure 5 Figure Supplement 1.**
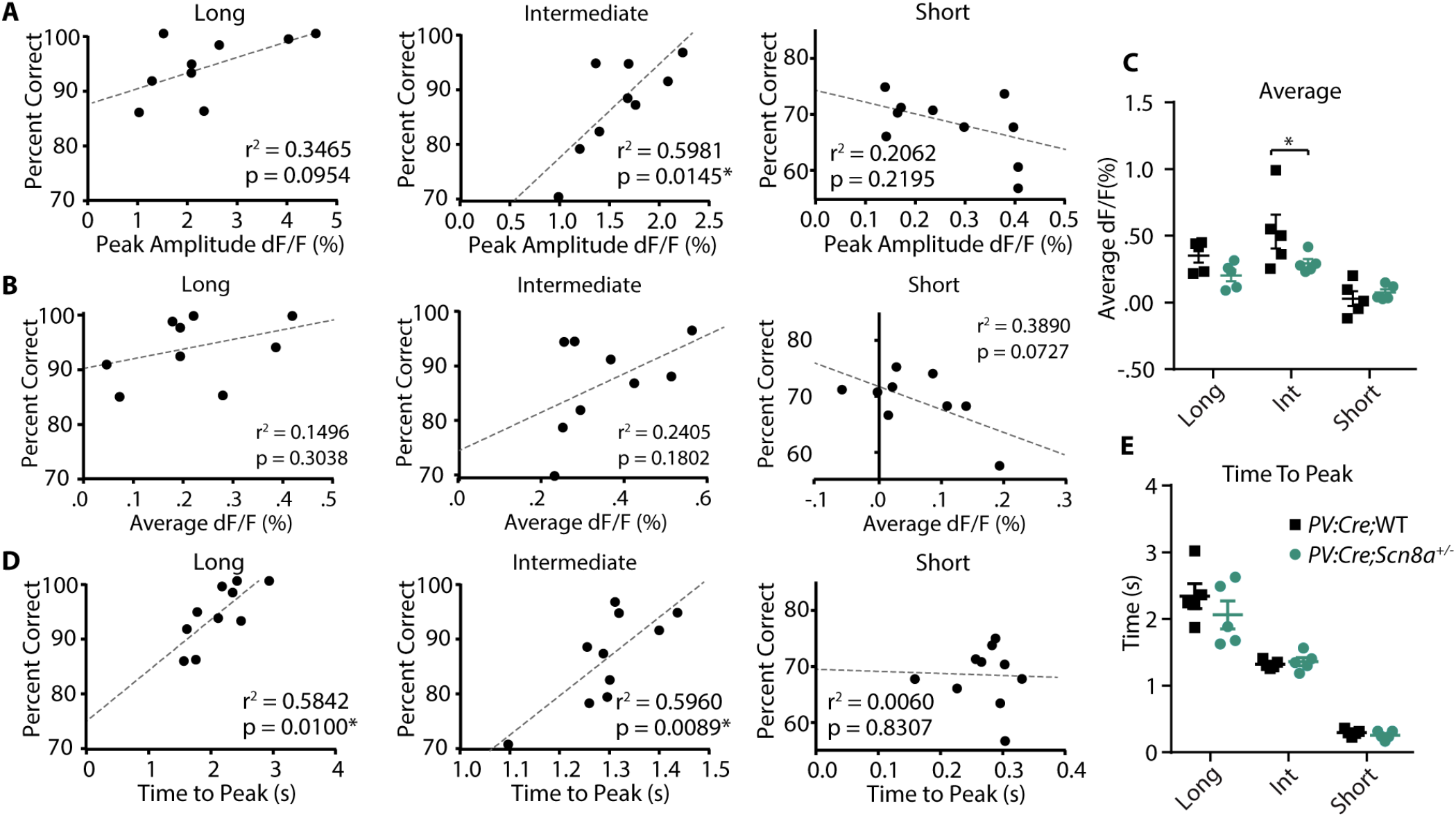
Peak Amplitude and Time to Peak of dF/F are predictors of performance at certain cue lengths, while average dF/F is not. Data points represent each animal’s average performance at each cue length plotted against the peak amplitude (A), average (B), and time to peak (D) and fit with a linear regression line. A, peak amplitude was a significant predictor of the percentage of correct choices for the int cue (r^2^ = 0.5981, p = 0.0145, n = 10 mice for all), but not at other cue lengths. B, Average dF/F did not predict performance at any cue length, but was reduced at the intermediate cue (p = 0.0466) for *PV:Cre;Scn8a*^+/−^ mice (C, two-way ANOVA, F _(1,24)_ = 4.658, p = 0.0411; n = 5 per group). D, Time to peak predicted performance at 5s and 2s (5s, r^2^ = 0.5842, p = 0.0100; 2s, r^2^ = 0.5960, p = 0.0089), E, However, time to peak is not different between groups (two-way ANOVA, F _(1, 24)_ = 0.9492, p = 0.3396).

**Figure 5 Figure Supplement 2.**
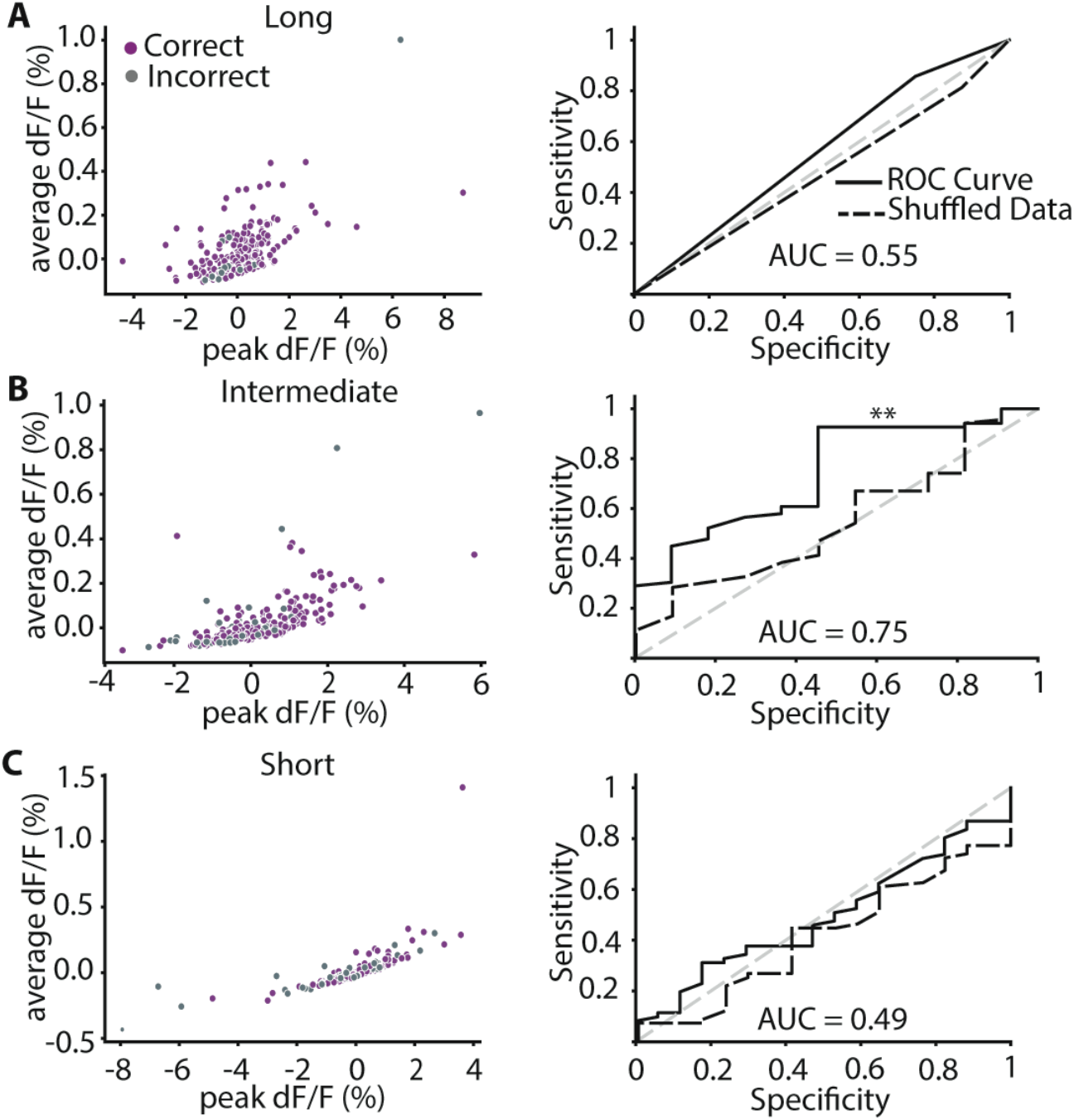
Trial Outcome can be predicted at the intermediate cue, but not the long or short cues. A-C (left), all trials for each cue length (long, intermediate, short) are plotted by their normalized peak and average dF/F, with correct trials in purple and incorrect trials in grey. Right, a random forest classifier trained on the features peak and time to peak, and then an ROC curve was generated (solid black line), and the AUC was calculated. For each cue length, the ROC curve based on the shuffled data was also generated (black dashed line). B, only for the intermediate cue was the classifier able to predict trial outcome at a level significantly greater than chance as shown in the ROC-AUC curve (AUC comparison, p = 0.0056).

**Figure 5 Figure Supplement 3.**
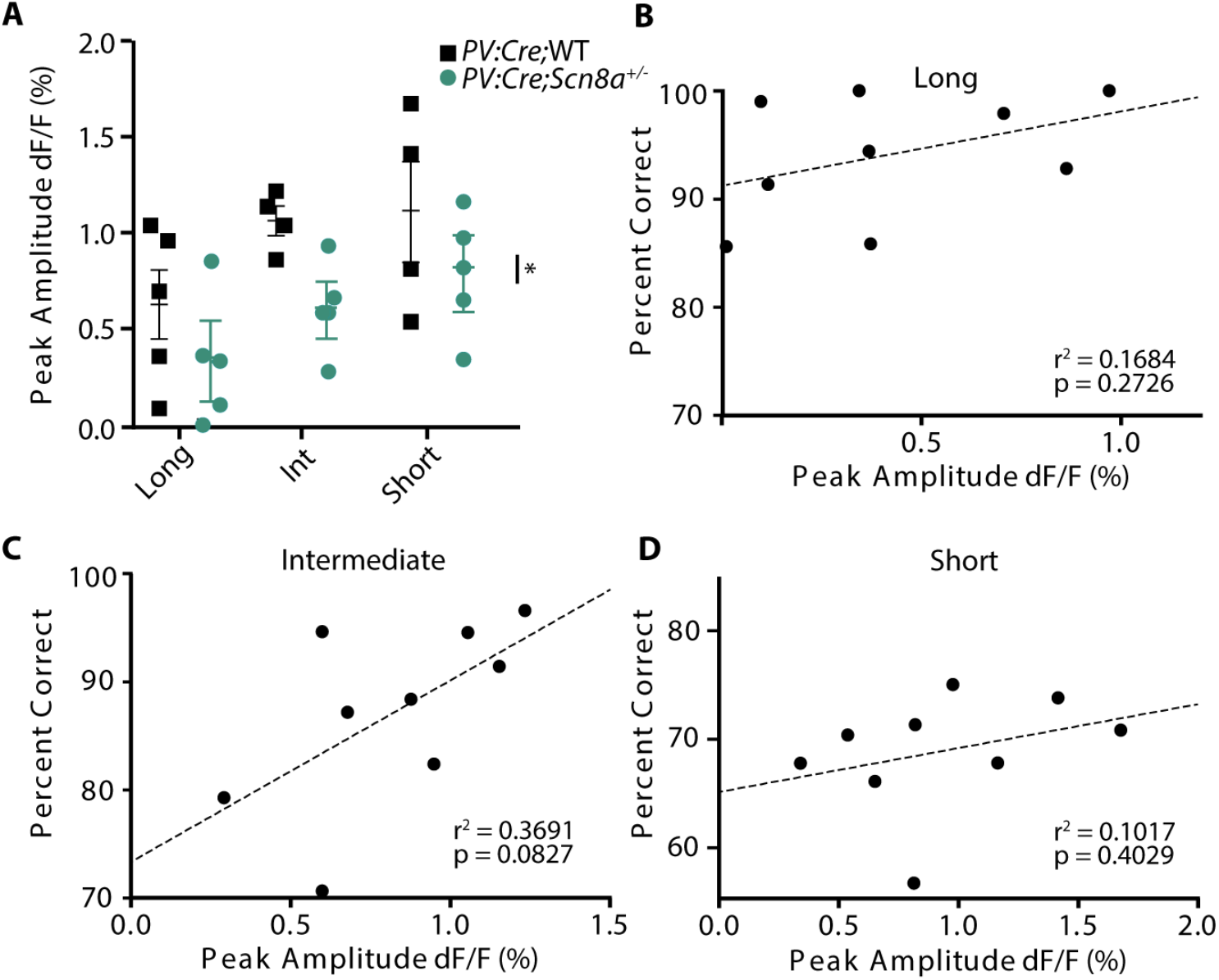
PVIN reward-related activity is reduced in Scn8a^+/−^ mice, but is not correlated to overall performance. A, *PV:Cre;Scn8a*^+/−^ mice exhibited decreased reward activity (two-way ANOVA, group effect, F _(1,25)_ = 8.13, p = 0.0086; n = 5 per group). B-D, Peak amplitude of reward activity was plotted against the average percent correct for each animal at each cue length. However, reward activity was not significantly correlated with overall performance, at any cue length. Data are shown as individual points or averages, and errors bars represent ± SE.

**Figure 7 Figure Supplement 1.**
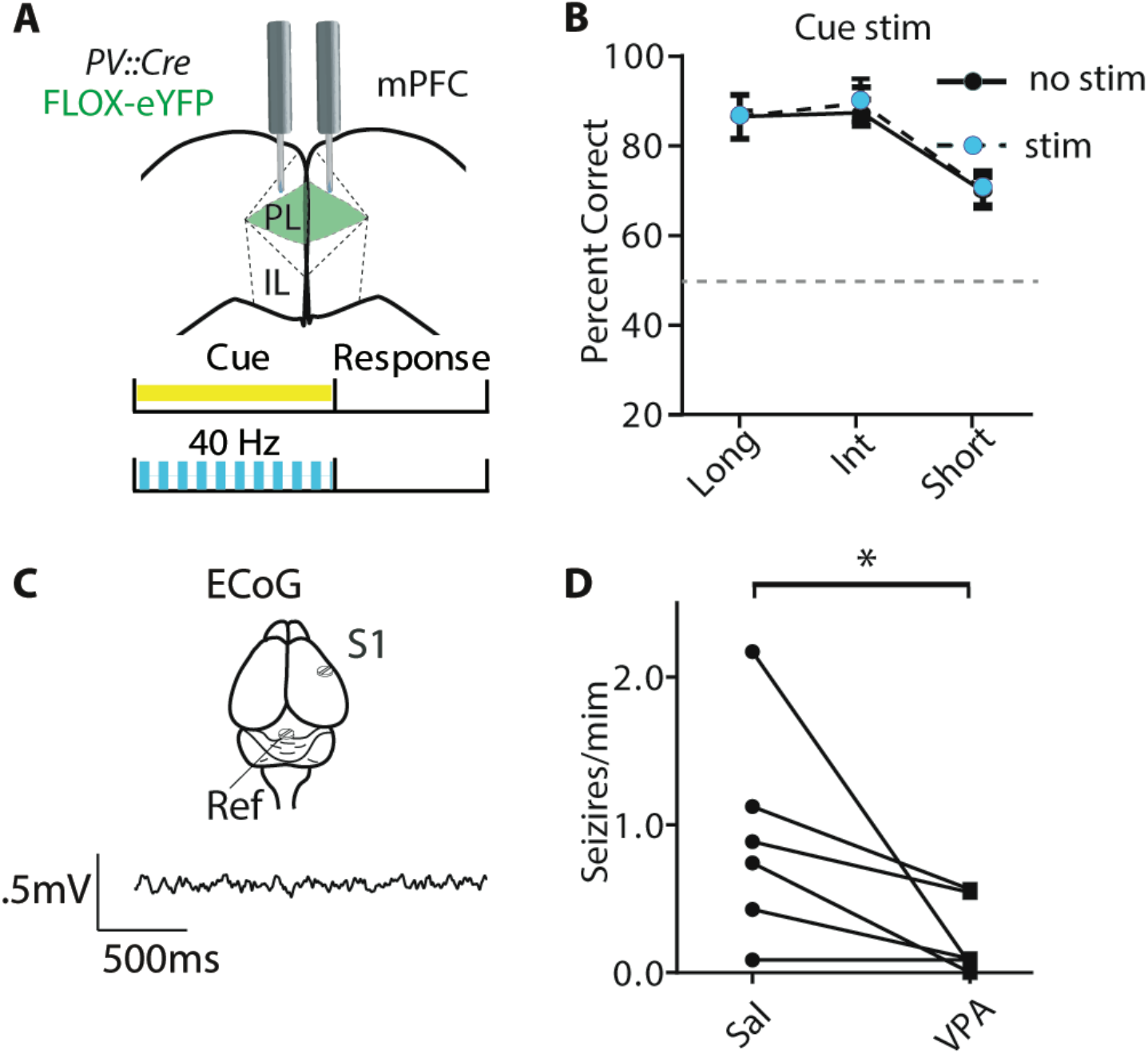
40 Hz stim has no effect on animals expressing the control virus and seizures are not significantly reduced by VPA. A, Mice received a bilateral injection with the FLOX-eYFP virus, and were implanted bilaterally with optical fibers for delivering blue light. B, There is no effect of light (two-way ANOVA, F _(1, 15)_ = .2598, p = 0.6177, n = 5 mice). C, Illustration of ECoG recording over S1 with reference screw in the cerebellum. D, VPA reduces the amount of seizures per minute (Wilcoxon matched-pairs signed rank test, p = 0.0313, n = 6).

## Source Data Files

Source data files for Figure 1, Figure 2, Figure 3, Figure 4, Figure 5, Figure 6, and Figure 7, along with Figure 2 - supplemental table 1, Figure 5 - Supplemental Figure 1-3, and Figure 7 - Supplemental Figure 1 are available at the following link: https://dataverse.harvard.edu/dataset.xhtml?persistentId=doi:10.7910/DVN/MCMHSW .

## Acknowledgements

BRF is supported by the Stanford School of Medicine Dean’s Fellowship, NIH-NINDS T32NS007280-35, NIH-NINDS Diversity Supplement to R01NS034774-24A1, and NIH-NINDS F32NS112764. JRH is supported by the NIH-NINDS R01NS034774-24A1 and NIH-NINDS T32NS007280-35. We thank Dr. Christina Kim for her consultation on the set up and implementation of the fiber photometry microscope and data analysis. We thank Dr. Ralf Wimmer for his consultation on the design and implementation of the behavior chamber and training paradigm for the attentional engagement task. We thank Dr. Wen-Jun Gao for his critical feedback on the manuscript. We thank Revathi Kaduru for her assistance in the electrocorticography experiments.

## Declaration of Interests

The authors declare no competing interests.

## Notes

### Competing Interest Statement

The authors have declared no competing interest.

